# Deciphering the Role of Aggrecan in Parvalbumin Interneurons: Unexpected Outcomes from a Conditional ACAN Knockout That Eliminates WFA+ Perineuronal Nets

**DOI:** 10.1101/2024.03.20.585910

**Authors:** Sverre Grødem, Elise H. Thompson, Malin B. Røe, Guro H. Vatne, Ingeborg Nymoen, Alessio Buccino, Tarjei Madland, Torkel Hafting, Marianne Fyhn, Kristian K. Lensjø

**Affiliations:** Department of Bioscience, University of Oslo, Norway; Allen Institute for Neural Dynamics, Seattle, USA; Institute for Basic Medical Research, University of Oslo, Norway

**Author notes:** indicates equal contribution (Sverre Grødem, Elise H. Thompson).

## Abstract

The transition from juvenile to adult is accompanied by the maturation of inhibitory parvalbumin-positive (PV+) neurons and reduced plasticity. This transition involves the formation of perineuronal nets (PNNs), a dense configuration of the extracellular matrix that predominantly envelops parvalbumin-positive (PV+) neurons. Aggrecan, a proteoglycan encoded by the ACAN gene, has been shown to have a key role in the PNNs as knock-out of ACAN in the adult brain reactivates juvenile plasticity, but the contribution of different cell populations is unknown

Here, we establish and characterize a mouse model in which ACAN is selectively knocked out (KO) in PV+ neurons (ACANflx/PVcre). Moreover, we develop a viral tool to perform similar cell-type directed KO in adult mice. Both models are compared with the traditional method of PNN removal, namely enzymatic degradation of PNNs with Chondroitinase ABC (chABC).

We show that PV+ neurons in adult ACANflx/PVcre mice do not produce PNNs that are labeled by *Wisteria floribunda agglutinin* (WFA), the most commonly used PNN marker. Surprisingly, electrophysiological properties of PV+ interneurons in the visual cortex (V1) and ocular dominance plasticity of adult ACANflx/PVcre mice were similar to controls. In contrast, AAV-mediated ACAN knockout in adult mice increased ocular dominance plasticity. Moreover, in vivo chABC treatment of KO mice resulted in reduced firing rate of PV+ cells and increased frequency of spontaneous excitatory postsynaptic currents (sEPSC), a phenotype associated with chABC treatment of WT animals. This suggests compensatory mechanisms in the germline KO. Indeed, qPCR of bulk tissue indicates that other PNN components are expressed at higher levels in the KO animals. Finally, we perform memory-and behavioral testing to see if the lack of ACAN from PV+ neurons throughout development affected stereotypic behaviors and memory processing. ACANflx/PVcre mice have learning and memory abilities similar to controls, but use bold search strategies during navigation in the Morris water maze. The low level of anxiety-related behavior is confirmed in an open field and zero maze, where they spent nearly twice the time in open areas.

## Introduction

Perineuronal nets (PNNs) are a specialized form of extracellular matrix (ECM) structures in the CNS involved in neural network maturation and regulation of brain plasticity. The PNNs constitute a dense mesh of components anchored to the cell membrane by hyaluronic acid synthetase, comprised of link proteins, tenascin-R, and chondroitin sulfate proteoglycans (CSPGs) such as versican, brevican, neurocan and aggrecan. Components of the PNN also bind to molecules that affect plasticity and function, such as Semaphorin 3A (Dick et al., 2013) and Otx2 (Beurdeley et al., 2012; Sugiyama et al., 2008). Aggrecan is a critical component for PNN assembly and structure, as pan-neuronal or virus-mediated knock-out of the ACAN gene prevents PNN formation or disrupts the structure, respectively (Rowlands, Lensjø, et al, 2018). Still, the contributions of the different PNN components and their production and regulation by different cell types remain elusive.

In the cerebral cortex, PNNs mostly enwrap basket-type Parvalbumin expressing (PV+) interneurons (Härtig et al., 1992; Lensjø, Christensen, et al., 2017). The PV+ neurons are potent regulators of cortical network function through their abundant connectivity with local excitatory and inhibitory neural networks (Hu et al., 2014). They innervate the soma and proximal dendrites of excitatory neurons, thereby strongly impacting spike timing and gain control in the cortex (Ferguson & Gao, 2018; Isaacson & Scanziani, 2011). In other areas, such as the amygdala, the hypothalamus, and CA2 of the hippocampus, PNNs also surround excitatory neurons (Morikawa et al., 2017). Both glial cells and neurons synthesize the major PNN components (Wiese et al., 2012), but merging evidence suggests that neurons can produce and regulate PNNs independently (Gottschling et al., 2019; Miyata et al., 2005; Rowlands et al., 2018).

In the mature brain, PNNs appear to influence PV+ interneuron function by restricting plasticity and facilitating the high-frequency firing activity of PV+ cells (Favuzzi et al., 2017; Lensjø et al., 2017; Marín, 2016; Tewari et al., 2018). How, and to which extent PNNs could facilitate high-frequency firing in PV+ neurons is unclear. Reports on PNN perturbation effects are contradictory; several patch clamp studies do not find any effect on firing frequency (Chu et al., 2018; Faini et al., 2018; Hayani et al., 2018), while some studies find significant reductions (Balmer, 2016; Favuzzi et al., 2017; Liu et al., 2022, 2023; Tewari et al., 2018). A recent review suggests that of the studies that have investigated PV cell firing rates following PNN perturbation, seven out of twelve studies found a decrease in firing rate, five out of twelve found no change, and zero out of twelve found an increase in firing rate (Wingert & Sorg, 2021). However, these studies are challenging to compare, as different PNN perturbations are applied to different brain regions and in mice of different ages. Moreover, the review pools extracellular and intracellular electrophysiology papers from in vivo and in vitro studies. The most commonly applied perturbation is enzymatic digestion of chondroitin sulfate glycosaminoglycans on CS-proteoglycans by chABC, but this is not specific to CSPGs in PNNs which only makes up 2-5% of the total CSPGs in the brain (Deepa et al., 2006). To add complexity, both the origin of the enzyme (e.g. Sigma or AMSBIO/Seikagaku), the concentration, volume, and incubation time are inconsistent between studies.

One paper that finds an effect on firing rate with chABC treatment suggests that an increased effective capacitance causes the reduced firing rate they observed in chABC-treated PV+ cells (Tewari et al., 2018). This theory is disputed by modeling the effect of increased capacitance on firing rate where they find that capacitance change alone cannot explain the large reduction (Hanssen et al., 2023). Other factors, like changes in reversal potentials and ion channel conductance, are suggested to be involved. One study that did not use chABC, focused on a transgenic or viral knockout of Brevican, a PNN proteoglycan, and found reduced firing rates in PV+ cells (Favuzzi et al., 2017). This paper did not look at effective capacitance, but did find a deficit in K_V_1.1 and K_V_3.1 potassium channels, which are important for the fast-spiking phenotype of PV+ cells (Du et al., 1996; Goldberg et al., 2008; Hanssen et al., 2023).

Ultimately, to determine if PNNs affect the fast-spiking phenotype of PV+ cells and synaptic plasticity, more precise perturbation is needed. To achieve this, we crossed a PVcre mouse (Hippenmeyer et al., 2005) with an ACAN-flox mouse (Rowlands et al., 2018), for targeted deletion of ACAN in PV+ cells. Knockout of ACAN from PV+ cells resulted in ablation of PNN formation on PV+ cells throughout the brain, indicating that ACAN expression in PV+ cells specifically is required for PNN formation. The knockout had no impact in areas where PNNs surround excitatory neurons, such as the hippocampal area CA2. Moreover, we find that germline KO of ACAN has only minute effects on the physiological properties of neurons and on memory processing, but our data indicates a change in phenotype with reduced anxiety compared to controls. In contrast, AAV-mediated genetic knockdown of ACAN in adult mice produced an elevated level of plasticity resulting in an ocular dominance shift in V1 after monocular deprivation, as was the case in the pan-neuronal ACAN KO (Rowlands et al., 2018), but no such shift was observed in the KO or controls. The discovery suggests that compensatory mechanisms might be in play in the ACANflx/PVcre, this theory is strengthened by our findings from qPCR of bulk tissue where we see that other PNN components are expressed at higher levels in the ACANflx/PVcre animals. Moreover, *in vivo* chABC treatment of adult KO mice reduced the firing rate of PV+ cells in V1, increased the frequency of sEPSCs, and reduced the amplitude of sIPSCs, similar to expected results after chABC treatment of WT mice. The lack of effects in the germline KO conflicts with current theories of the function of PNNs enwrapping PV+ neurons, and raises concerns with the correlation between chABC treatment effects and PNN perturbation.

## Results

### PV+ neuron targeted knock-out of ACAN prevents formation of PNNs

To achieve specific knockout of PNNs in PV+ interneurons, we crossed a cre-conditional ACANflox line with a PVcre driver line (ACANflx/PVcre). The PVcre line also served as the control group; cre expression is not expected to influence Aggrecan expression (Devienne et al., 2021; Favuzzi et al., 2017; Hippenmeyer et al., 2005). We mainly focused our investigation on the primary visual cortex (V1), a brain area where PNNs are believed to be involved in the regulation of postnatal plasticity.

Immunostaining for Aggrecan, PV and Wisteria Floribunda Agglutinin (WFA), a lectin commonly used to label PNNs, demonstrated that aggrecan and WFA were abolished in PV+ cells in the PV-selective ACAN KO (Fig. 1 and S1). Immunohistochemistry revealed that non-PV neurons were still sparsely encapsulated in PNNs; this was detected with both WFA and aggrecan staining (Fig. 1 and S1). The CA2 of the hippocampus is particularly noticeable in the ACANflx/PVCre line (Fig. 1C and D) because here PNNs are formed around excitatory neurons (Carstens et al., 2016; Lensjø, Christensen, et al., 2017). These results indicate that PNN formation around PV+ is dependent on their own expression of ACAN.

**Figure 1.**
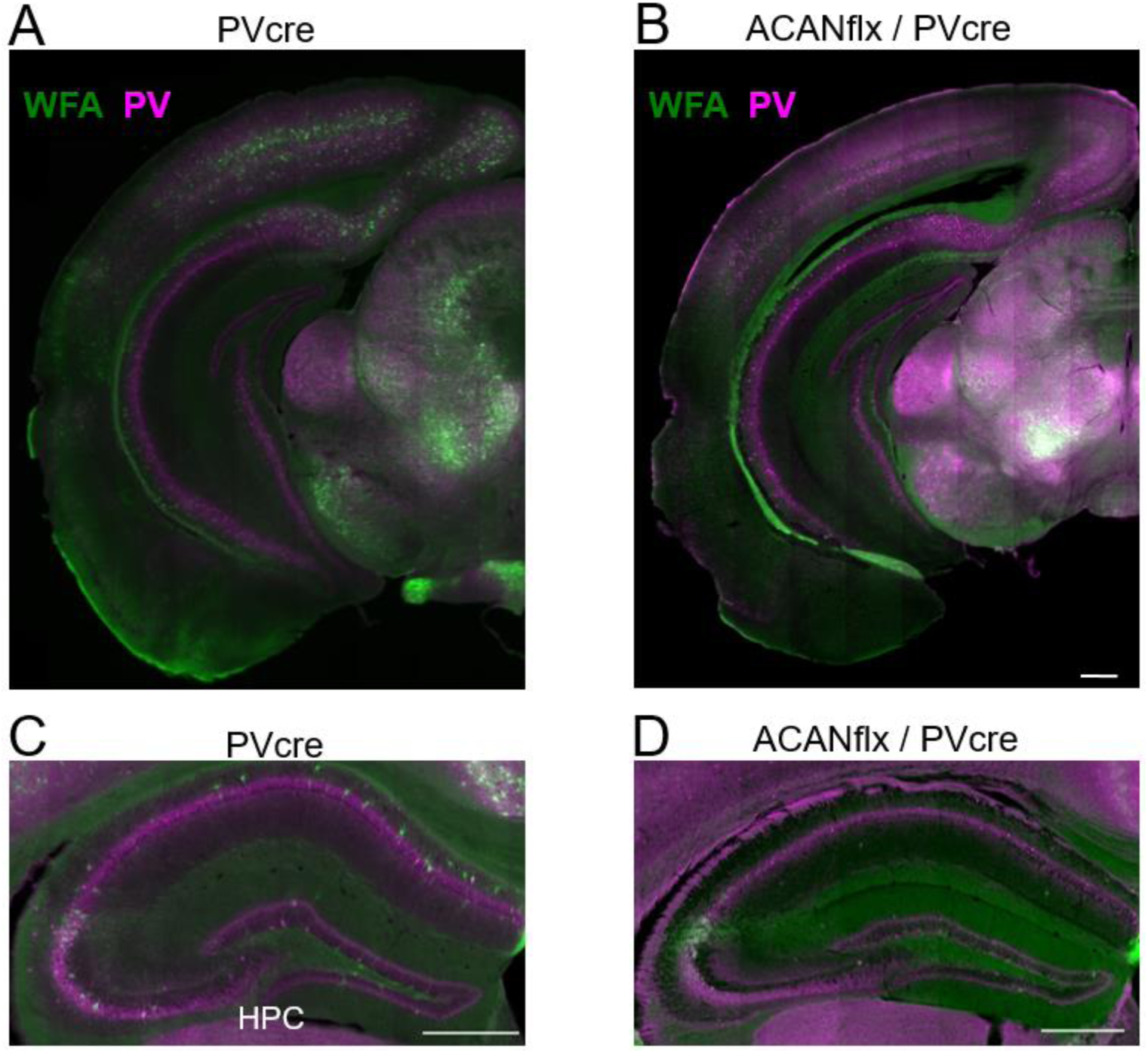
Acan knockout from PV+ cells (PVcre/Acan-loxP cross) causes a brain-wide depletion of WFA+ PNNs around PV+ neurons. **A)** Coronal section from a PVcre mouse stained with Wisteria Floribunda agglutinin (WFA) (green) and PV (magenta). **B)** Coronal section from a ACANflx/Pvcre mouse stained with WFA (green) and PV (magenta). **C)** Coronal section showing the hippocampus (HPC) of a PVcre mouse stained with WFA (green) and PV (magenta). **D)** Coronal section showing the hippocampus (HPC) of a ACANflx/PVcre mouse stained WFA (green) and PV (magenta). Scale bar = 500 µm

We next hypothesized that when aggrecan is knocked out, other PNN or ECM components may be upregulated as a potential compensatory mechanism. Accordingly, we investigated the expression in V1 of the PNN components neurocan, and tenascin-R, in addition to aggrecan (Fig. 2A). As a first and coarse measure, we used immunohistochemistry, but did not observe large-scale changes in the expression of neither neurocan and tenascin-r in the absence of aggrecan in PV+ cells (n = 3 ACANflx/PVcre or PVcre mice per staining group, six slices per animal). However, in qPCR analysis of gene expression in bulk tissue from V1, we observed divergent effects on gene expression in the KO vs control, and in an adult virus-mediated KO (Fig. 2B). To achieve KO in adults, we used retro-orbital injections of AAV-PV:mGreenLantern-P2A-cre virus, with a PV enhancer (Vormstein-Schneider et al., 2020) to target expression of Cre to PV+ neurons across the brain. qPCR analysis revealed that while ACAN expression is reduced in the germline KO, it is not eliminated, suggesting that while PV+ production of ACAN is required for WFA+ PNNs, ACAN produced by PV cells only represents a fraction of the total ACAN in the tissue (Fig. 2B). The ACANflx/PVcre mice also displayed higher neurocan and tenascin-r expression, and reduced expression of semaphorin-3a (Fig. 2B). The PV expression was below the qPCR detection limit for both the PVcre control line, and the adult KO, but was abundant in the virus-mediated adult KO (Fig. 2B). Notably, this stark difference in PV expression is not apparent in immunohistochemistry (Fig.2A)

**Figure 2.**
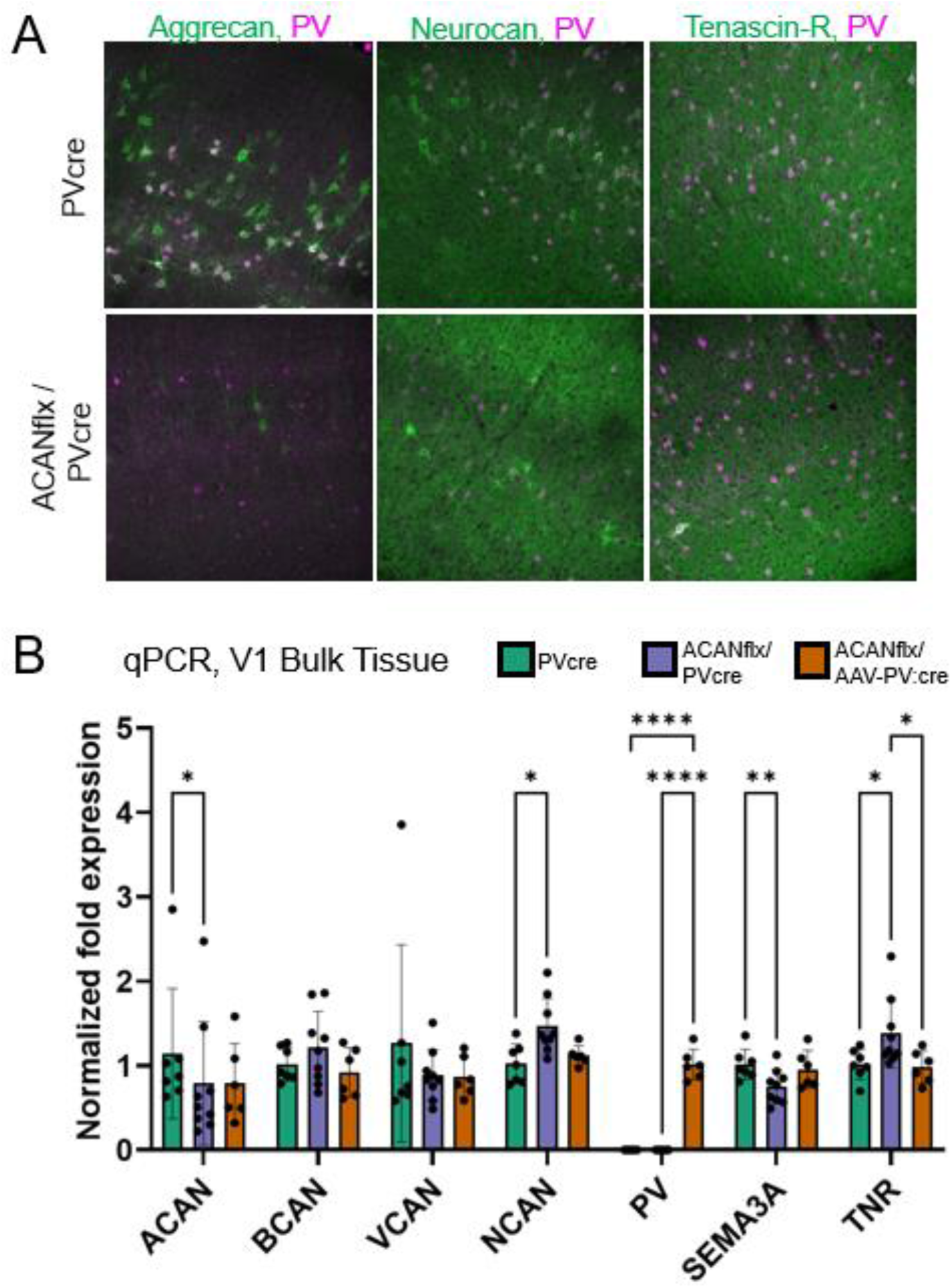
Immunohistochemistry and qPCR on V1 tissue reveals divergent effects on PNN gene expression in germline vs adult KO. **A)** Immunostaining for aggrecan, neurocan, tenascin-r and PV in ACANflx/PVcre and PVcre mice V1. **B)** qPCR on bulk tissue from V1 indicated altered gene expression in the germline knockout (ACANflx/PVcre) and adult knockout (ACANflx/AAV-PV:cre) compared to controls (PVcre). ACAN expression in bulk tissue was reduced but not eliminated in the germline knockout. NCAN and TNR gene expression was increased in the germline knockout, whereas SEMA3A expression was reduced. Gene expression in the adult knockout was only significantly different from control for PV, which was below the detection limit for the control and the germline KO. All samples were ran in duplicates or triplicates and averaged before further analysis. Tukey’s multiple comparisons test was performed. Bars represent mean ± SD, each dot represents one animal.

### Germline disruption of WFA+ PNNs does not affect intrinsic electrophysiology of PV+ neurons

To investigate if a knockout of ACAN in PV+ cells also influenced intrinsic properties of neurons, we obtained whole-cell patch clamp recordings from fast-spiking PV+ neurons from V1 in acute slices from adult ACANflx/PVcre mice, and PVcre control mice. The PV+ cells were labeled using a systemic Cre-dependent Flex-TdTomato PHP.eB AAV, which was injected intravenously at least three weeks before experiments. The cell type was confirmed based on fast firing properties (>200 Hz), and little to no spike adaptation.

These recordings revealed no differences between PV+ cells in the control and PV+ cells lacking aggrecan (Fig. 3A-H). No difference was found in resting membrane potential (Fig. 3H), input resistance or FI curves (Fig. 3E), despite the absence of WFA+ PNNs on the targeted PV+ neurons. Nor did we find any change in effective capacitance, which has been posited to be the cause of reduced firing rate in PNN deficient PV+ cells (Tewari et al., 2018).

**Figure 3.**
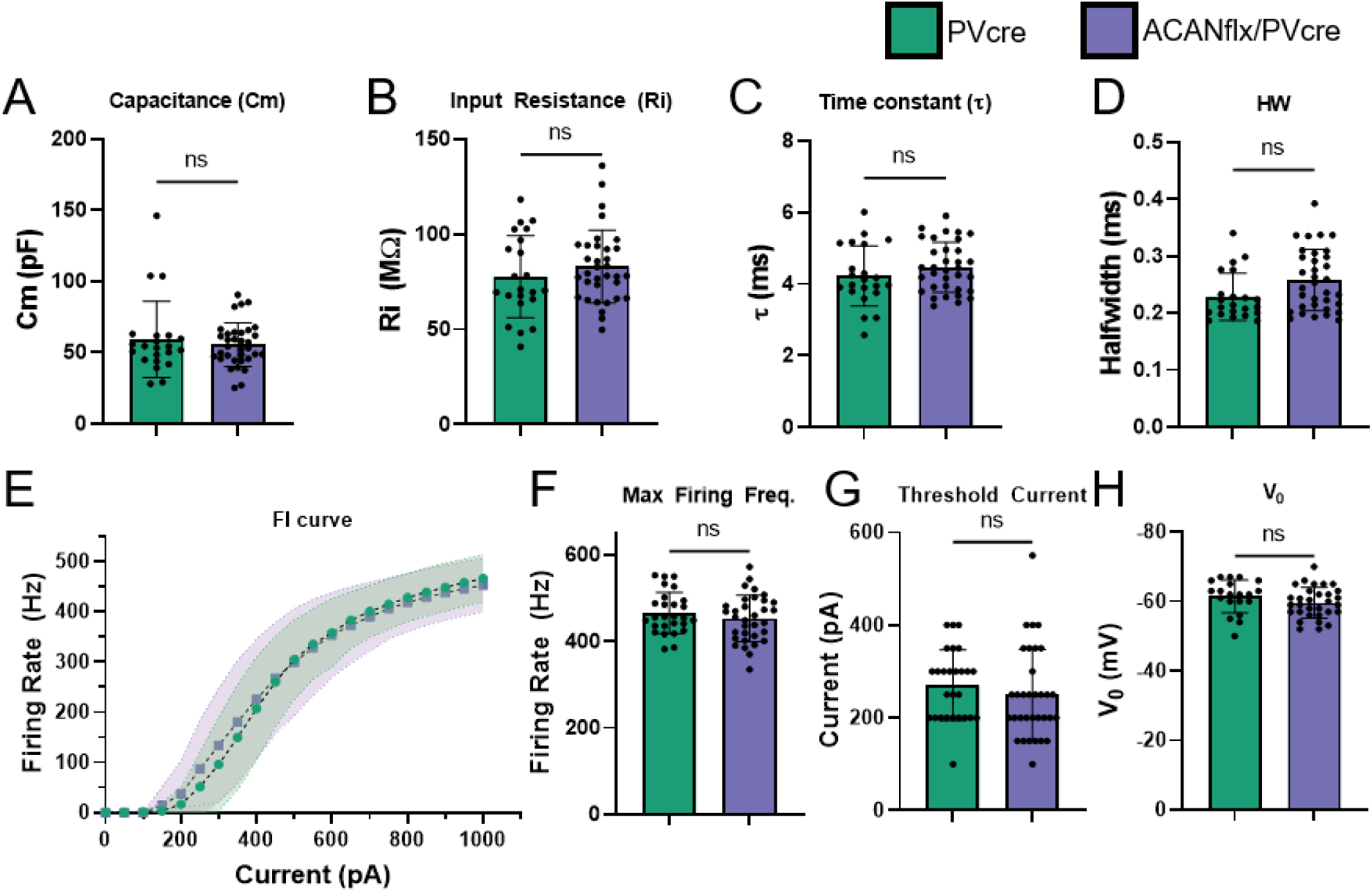
Intrinsic electrophysiological and firing properties of PV neurons are not affected by PV-specific germline KO of ACAN. **A)** Capacitance (pF). **B)** Input Resistance (MΩ). **C)** Time Constant (ms) **D)** Action potential half width (ms). **E)** FI curve, firing rate (Hz) as a function of injected current (pA). **E)** Maximum firing frequency (Hz). **F)** Maximum firing frequency (Hz). **G)** AP threshold current (50pA increments, pA). **H)** Resting membrane potential (mV). Each dot represents one cell. PVcre (green) n = 22/13 (cells/mice), AcanKO (blue) n = 34/14 for A, B, C, D; n = 27/14 and 33/14 for E, F, G; n = 20/12 and 31/12 for H. Data shown as mean ± SD. Unpaired student’s t-test, ns: p > 0.05.

### PV+ specific deletion of ACAN in adult mice, but not germline deletion, causes an ocular dominance shift after monocular deprivation

The synchronized activity of PV+ interneurons is thought to drive theta and high gamma oscillations and inhibition of these neurons suppresses this oscillatory activity (Amilhon et al., 2015; Cardin et al., 2009; Sohal et al., 2009). Emerging evidence suggests that PNN removal causes a reduction in PV+ interneuron signaling (Favuzzi et al., 2017; Lensjø et al., 2017; Tewari et al., 2018). Because the literature suggests the presence of a strong connection between the intrinsic properties of PV+ interneurons and the presence of PNNs, it was surprising that our *ex vivo* recordings revealed no effect of ACAN KO in PV+ cells. Nevertheless, developmental effects caused by obstructed PNN formation could have influenced the network activity. Therefore, we recorded the local field potential *in vivo* during spontaneous and evoked responses in the binocular zone of the primary visual cortex (V1B) (Fig 4A). During critical period plasticity in the visual cortex, the area is highly susceptible to changes in activity. If one eye is deprived of input, even for a short period, it creates an ocular dominance shift in the V1B towards the open eye (Frenkel & Bear, 2004; Gordon & Stryker, 1996; Wiesel & Hubel, 1963). In addition to affecting PV+ neuron properties, PNN assembly marks the closure of critical periods. When PNNs are removed in an adult, it re-opens the critical phase of plasticity (Lensjø et al., 2017; Pizzorusso et al., 2002; Rowlands & Lensjø et al., 2018).

**Figure 4:**
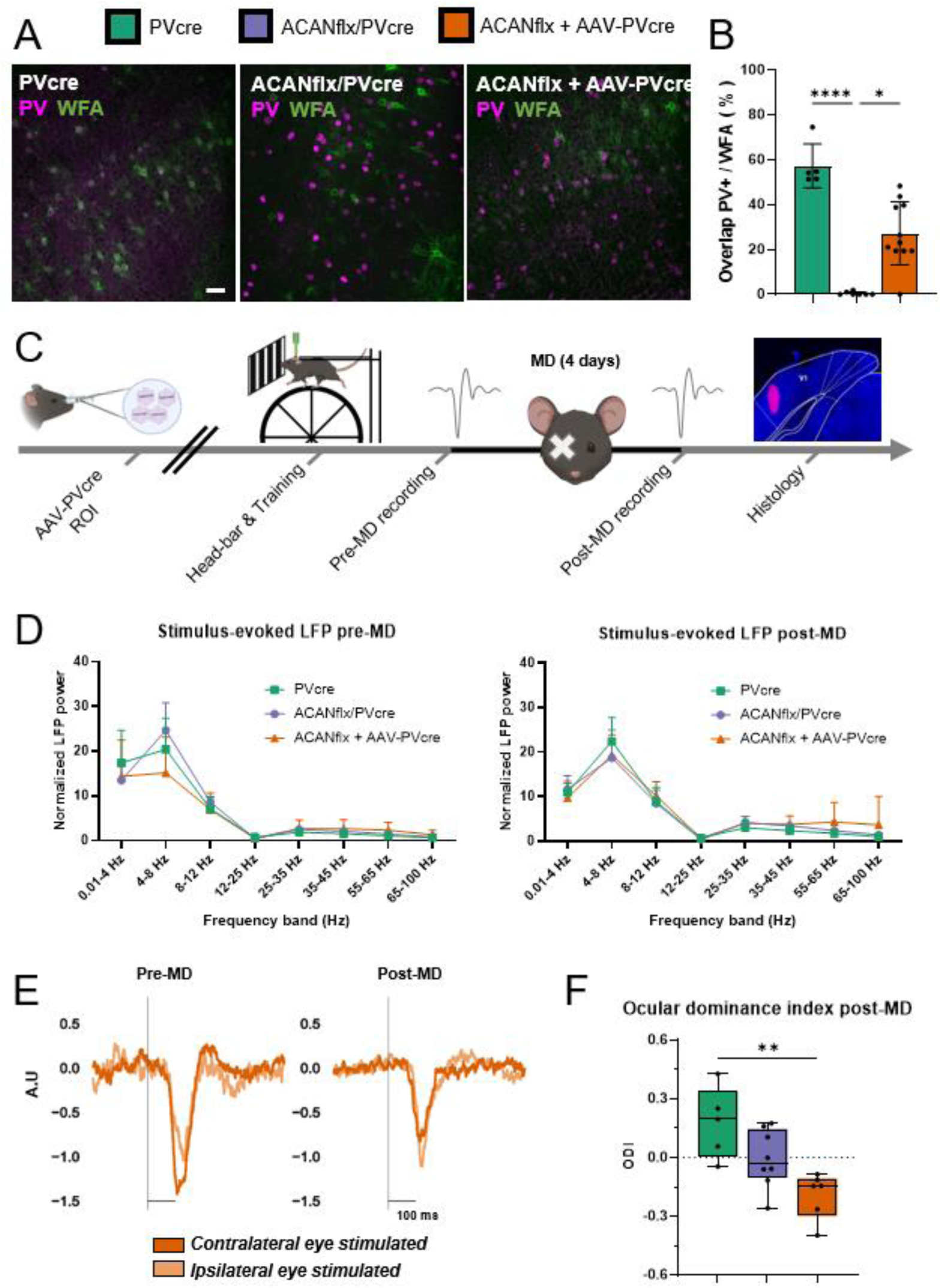
Network properties and ocular dominance plasticity; PV+ specific deletion of aggrecan from adult, but not germline, causes an ocular dominance (OD) shift after monocular deprivation (MD). A) Timeline for ocular dominance plasticity experiments. AAV-PVcre virus was delivered in ACANflx mice with a retroorbital injection (ROI) two months before headbar implantation. Headbar implantation was the first procedure for PVcre and ACANflx/PVcre mice before all groups were trained on a running wheel prior to recording extracellular activity. Local field potentials (LFP) were recorded before and after 4 days of monocular deprivation (MD). The silicone probe was coated in DiI on the last day of recording, histological verification of probe position was done post mortem as shown in the picture (far-right panel). B) PNNs visualized using WFA staining (green) together with PV labeling (magenta) in V1 from PVcre (left), ACANflx/PVcre (middle) and ACANflx+AAV-PVcre (right). Scale bar=50µm. C) Efficiency of AAV-PVcre virus transfection was found by quantifying PV/WFA overlap relative to total number of PV+ neurons (%). ACANflx + AAV-PVcre mice were different to PVcre (p-value 0.0013) and ACANflx/PVcre (p-value 0.0210). ACANflx/PVcre mice were also different to PVcre mice (p-value 0.0012). PVcre n=6, ACANflx/PVcre n=7, ACANflx + AAV-PVcre n=8, Mann-Whitney test used. D) Stimulus evoked normalized LFP power for different frequency bands (Hz) before and after MD. Pre-MD: ACANflx/PVcre mice were different from PVcre for 0.1-4 Hz (p-value 0.0056) and from ACANflx + AAV-PVcre for 4-8 Hz (p-value 0.0008). PVcre n=6, ACANflx/PVcre n=5, ACANflx + AAV-PVcre n=4. Post-MD: ACANflx/PVcre and ACANflx+AAV-Pvcre were significantly different from PVcre for 4-8 Hz (p-values 0.0053 and 0.0035, respectively). PVcre n=6, ACANflx/PVcre n=8, ACANflx + AAV-PVcre n=5. 2-way ANOVA with Tukey’s multiple comparison test used. E) Average normalized LFP responses of all ACANflx+AAV-PVcre mice (PreMD n=4, PostMD n=6) showing an ocular dominance shift after MD. Ocular dominance index (ODI) was calculated from normalized LFP responses (A.U) to visual stimulation of the ipsi- and contralateral eye before and after monocular deprivation (MD). F) Ocular dominance shift was observed in ACANflx+AAV-PVcre (p-value 0.0439). There was also a difference between PVcre and ACANflx+AAV-PVcre mice (p-value 0.0008) post-MD. Pre-MD: PVcre n=6, ACANflx/PVcre n=5, ACANflx + AAV-PVcre n=4. Post-MD: PVcre n=5, ACANflx/PVcre n=8, ACANflx + AAV-PVcre n=6. ANOVA with Tukey’s multiple comparisons test.

We sought to illuminate whether the PVcre/ACANflx mice, that never had WFA+ PNNs on PV+ cells, had a life-long state of high plasticity. We reasoned that despite lacking PNNs, compensatory mechanisms during development could have ensured the maturation of the PV+ neurons and closure of the critical period. To circumvent potential developmental effects, we also sought to perform the experiment with animals where aggrecan was knocked out in PV+ cells in adult mice using a systemic AAV expressing Cre under a PV enhancer. The efficacy of the virus transduction was determined by quantifying PV/WFA overlap relative to the total number of PV+ neurons. ACANflx/AAV-PVcre mice had an intermediate amount of WFA/PV overlap compared to PVcre and the ACANflx/PVcre cross (Fig. 4B-C, PVcre: 63.64% ± 10.31%, ACANflx + AAV-PVcre: 28.99% ± 21.31%, ACANflx/PVcre: 0.41% ± 0.74%, based on images from V1). As anticipated, the largest difference was found when comparing PVcre and ACANflx/PVcre (Fig. 4C, p-values PVcre vs. ACANflx/PVcre: 0.0012, PVcre vs. ACANflx+AAV-PVcre: 0.0013, ACANflx/PVcre vs. ACANflx+AAV-PVcre: 0.021. Mann-Whitney test). The PV/WFA overlap in PVcre control animals (63.64% ± 9.41%) is in agreement with a recent, extensive study on the distribution of PNNs in the mouse brain (Lupori et al., 2023).

Having confirmed that adult knock-out of ACAN attenuated PNN expression, we conducted recordings to assess stimulus-evoked LFP activity and investigate ocular dominance plasticity after MD. Using acutely inserted 32-channel silicon probes, we recorded extracellular activity during periods of spontaneous activity and during visual stimulation (100% contrast sinusoidal gratings) while covering the contralateral, the ipsilateral or neither of the eyes. Activity was recorded before and after monocular deprivation (MD). Animals were head-fixed and ran on a running wheel during the experiment. The ocular dominance index (ODI) was calculated based on the visually evoked local field potential (LFP) recorded during stimulation of the contralateral and ipsilateral eye independently (ODI = contralateral - ipsilateral/contralateral + ipsilateral).

Our results exhibited no significant differences between ACANflx/PVcre, ACANflox + AAV-PV:cre, or PVcre controls in LFP activity evoked by visual stimulation in any frequency band (Fig. 4F, 0.01-4, 4-8, 8-12, 12-25, 25-35, 35-45, 55-65 and 65-100 Hz). These findings indicate that a reduction of WFA+ PNNs in the virus group did not have an impact on network activity *in vivo* (Fig 4D). Surprisingly, no significant shift in the ODI was detected in the ACANflx/PVcre group, suggesting that their state of plasticity was comparable to controls (Fig. 4H, p-values ODI pre- vs post-MD PVcre: 0.0966, ACANflx/PVcre: 0.500, unpaired t-test). However, recordings of the visually evoked potentials before and after MD revealed a shift in ocular dominance towards the ipsilateral eye in the ACANflx/AAV-PV- cre mice (Fig. 4E and F, p-value 0.0096 pre- vs post-MD, unpaired t-test). Whether the PNN modification caused an opening of a “juvenile-like” critical period plasticity or an elevated level of adult-type plasticity cannot be concluded from our data. It should be noted that investigations of ocular dominance shifts are commonly performed in anesthetized animals, while we recorded from awake animals in which the neural activity is different from that of anesthetized animals (Aasebø et al., 2017).

### ACANflx/PVcre and PVcre mice employ different strategies during spatial learning

We used the Morris water maze to study learning, memory and cognitive flexibility in ACANflx/PVcre mice (Morris, 1981). The water maze is a navigation task where an animal is placed in a circular pool to search for a platform hidden below the surface. The water is made opaque so that the platform is not visible, and the animals must learn to swim to the platform from four different starting points along the walls of the pool. The training consisted of four daily trials for seven days, and performance was scored as average daily path length and latency to platform. Both PVcre and ACANflx/PVcre mice learned the task within this seven-day period (Fig. 5A and B, row factor (days of training), p-value < 0.0001, repeated measures ANOVA). However, ACANflx/PVcre mice had shorter path length and latency to platform over the course of training compared to PVcre mice (Fig. 5C and D, mean path length (cm) PVcre 509.7 cm ± 48.30, ACANflx/PVcre 394.4 cm ± 65.96, p-value = 0.002. Mean latency (s) PVcre 33.25 s ± 5.14, ACANflx/PVcre 22.9 s ± 6.72, p-value = 0.006, unpaired Student’s t-test). General swimming abilities, such as swim speed, were comparable to controls (Day 1 mean speed PVcre: 0.12 ± 0.02, ACANflx/PVcre: 0.1 ± 0.03, p-value: 0.11, unpaired Student’s t-test, Fig. S4A). To investigate why the ACANflx/PVcre mice performed better during training, we analyzed different behavioral features of the training. We defined six swimming patterns based on previous studies: Thigmotaxis, Random search, Scanning, Indirect search, Directed search and Direct path (Fig. 5E) (Cooke et al., 2019; Illouz et al., 2016). Some swimming patterns have a higher probability of finding the platform and are scored accordingly. This is called a “cognitive score”, where the highest score is assigned to a swimming pattern aimed directly at the platform.

**Figure 5.**
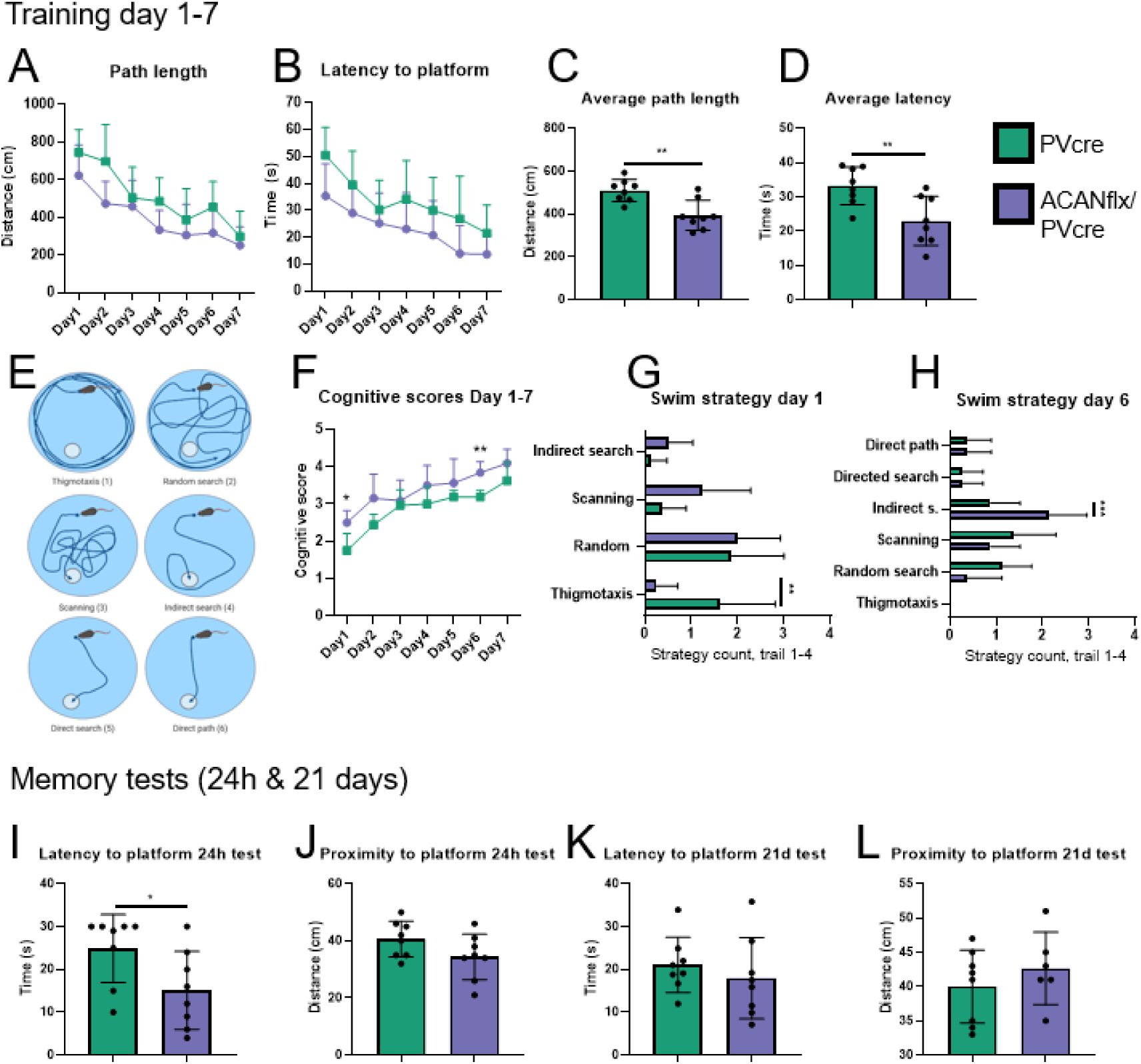
ACANflx/PVcre and PVcre mice use different strategies in the Morris water maze, potentially due to lowered anxiety in mice lacking WFA+ PNNs in PV+ cells. A) Swimming path length (cm) during training day 1-7. Both groups showed decreasing swim path length from day 1-7 (Row factor (days of training), p-value < 0.0001). B) Latency to platform (s) during training day 1-7 (Row factor (days of training), p-value < 0.0001). C) Path length (cm) averaged over all days of training (mean path length (cm) PVcre 509.7 cm ± 48.30, ACANflx/PVcre 394.4 cm ± 65.96, p-value = 0.002). D) Latency (s) averaged over all days of training (mean latency (s) PVcre 33.25 s ± 5.14, ACANflx/PVcre 22.9 s ± 6.72, p-value = 0.006). E) Swim strategies and associated cognitive scores. Thigmotaxis (1), random search (2), scanning (3), indirect search (5), direct search (4), direct path (6). F) Cognitive scores given from day 1-7 of training. Cognitive score on day 1 and day 6 of training differed between ACANflx/PVcre and PVcre (cognitive score day one: PVcre 1.75 ± 0.43, ACANflx/PVcre 2.5 ± 0.31, p-value = 0.02; day six: PVcre 3.19 ± 0.16, ACANflx/PVcre 3.84 ± 0.28, p-value = 0.0014). G) Swim strategy day 1 (mean prevalence of thigmotaxis PVcre 1.62 ± 1.11, ACANflx/PVcre 0.25 ± 0.43, p-value = 0.006). H) Swimming strategy day 6 (mean prevalence of indirect search PVcre 0.88 ± 0.59, ACANflx/PVcre 2.13 ± 0.78, p-value = 0.0004). I) Latency (s) to platform zone during memory test 24 hours after training (mean latency (s) PVcre 24.88 s ± 7.42, ACANflx/PVcre 15.13 s ± 8.50, p-value = 0.04). J) Proximity (cm) to platform during the 24 hour memory test. K) Latency (s) to platform zone during memory test 21 days after training L) Proximity (cm) to platform during 21 day test. Repeated measures ANOVA with Sidak’s multiple comparisons test used for A, B, F, G, H. Unpaired Student’s t-test used for C, D and I-L. Data shown as mean ± SD. Each dot represents one animal. PVcre (green) n= 8, ACANflx/PVcre (purple) n= 8.

Early in the training, rodents utilize “scanning” and “random search” behaviors to find the platform. During random search, the animal swims along the edges as well as the rest of the pool. During scanning, the mouse largely avoids the edges. Because the platform is situated away from the edge of the pool, there is a higher chance of finding the platform using the “scanning” strategy. We found that ACANflx/PVcre mice tended to rely on scanning more frequently compared to PVcre controls, which resulted in them finding the platform on the first trial, on day one (Fig. 5G). The occurrence of thigmotaxis, a sign of anxiety in rodents where they stay close to the edges of their environment (Treit & Fundytus, 1988), was very low among ACANflx/PVcre (Fig. 5F and G, PVcre 1.62 ± 1.11, ACANflx/PVcre 0.25 ± 0.43, p-value = 0.006, repeated measures (RM) ANOVA with Sidak’s multiple comparisons test). The ACANflx/PVcre group demonstrated superior strategy on day one and six, reflected in higher cognitive scores compared to PVcre controls (Fig. 5H, cognitive score day one: PVcre 1.75 ± 0.43, ACANflx/PVcre 2.50 ± 0.31, p-value = 0.02; day six: PVcre 3.19 ± 0.16, ACANflx/PVcre 3.84 ± 0.28, p-value = 0.0014). Results from the training indicate that the ACANflx/PVcre mice’s superior performance is due to a lower level of anxiety related behavior.

To test the ability of the mice to remember the platform location, we performed two probe trials where the platform is removed and initial latency to the platform zone and mean proximity to the platform zone throughout the test is used as an indicator of memory. One test was done after 24 hours, and the next after 21 days. In the 24-hour probe trial, latency to the platform zone was shorter in the ACANflx/PVcre group (Fig. 5I, PVcre 24.88 s ± 7.42, ACANflx/PVcre 15.13 s ± 8.50, p-value = 0.04, unpaired Student’s t-test), perhaps suggesting that the ACANflx/PVcre were quicker to choose the correct swim path leading to the platform zone. Their memory of the platform location was similar to controls, reflected in proximity to the platform throughout the probe trial (Fig. 5J). The 21-day memory test did not reveal any significant differences between the groups (Fig. 5K and L).

To test whether anxiety affected learning, we re-trained the animals with a new platform location, so-called reversal learning. The animals are now habituated to the water maze and anxiety should not have an impact on learning. Reversal learning also reflects cognitive flexibility, an ability that relates to level of plasticity (Happel et al., 2014). During reversal training, we found no differences in learning abilities (Fig S4A and B). It was evident that the mice knew which swim strategies to utilize to succeed. On day one of the original training, all mice used the “random search” strategy approximately 50% of the time; while on day one of the reversal training, it had decreased to approximately 25% (Fig. 5G and S4D). In the PVcre group, the amount of “scanning” behavior had increased from 9% on day one of the original training, to 56 % on day one of the reversal training. The ACANflx/PVcre mice kept a shorter proximity to the platform location during the 24-hour test (Fig. S4F, PVcre 54.63 cm ± 9.9, ACANflx/PVcre 41.13 cm ± 8.3, p-value = 0.02, unpaired Student’s t-test) while latency (s) to platform was similar (Fig. S4E). No significant differences were detected in the 21-day test (Fig. S4G and H).

Overall, our results indicate that ACANflx/PVcre mice have a slightly better ability to learn and remember the Morris water maze task. The low occurrence of thigmotaxis in the initial training round and the use of scanning early in training seem to give the ACANflx/PVcre mice an advantage. This might be caused by a low level of anxiety and/or reduced risk assessment. These results reflect the shortcomings of the water maze task, where animals with low anxiety perform better regardless of cognitive abilities (Darcet et al., 2014; Higaki et al., 2018; Pritchett et al., 2016).

### ACANflx/PVcre mice have lower levels of anxiety-like behavior

The results from the water maze indicated that the ACANflx/PVcre mice had a lower level of anxiety. Because mice are prey animals, they will naturally spend more time along edges and avoid open spaces where they risk being spotted by predators. Their explorative nature drives them to explore their environment in short running bouts across the open space. To test their performance in an open environment, the mice were placed in the center of an open field box (50×50×50 cm box) and then left to explore it for 5 minutes (Fig. 6).

**Figure 6.**
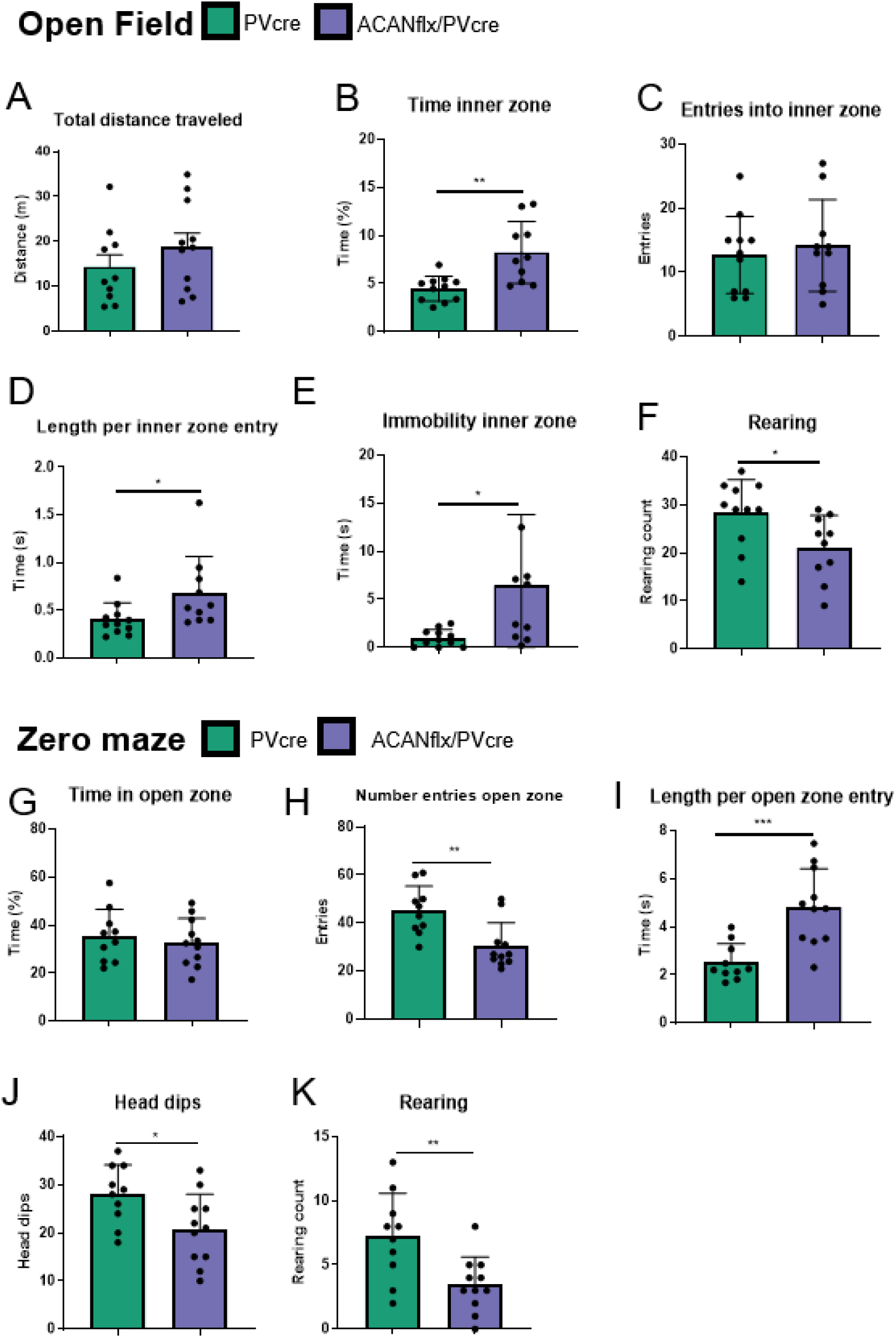
AcanKO demonstrates a lower level of anxiety and risk assessment behavior in the open field and zero maze compared to controls. Open field. **A)** Total distance traveled (m) in the open field. **B)** Percentage of time spent in the inner zone of the open field (PVcre 4.47% ± 1.25, AcanKO 8.24% ± 3.1, p-value = 0.002). **C)** Total number of entries into the inner zone of the open field. **D)** Average time (s) spent in the inner zone per entry (PVcre 0.39 s ± 0.17, AcanKO 0.68 s ± 0.36, p-value = 0.04). **E)** Immobility (s) in the inner zone (PVcre 0.98 s ± 0.85, AcanKO 6.44 s ± 6.99, p-value = 0.02), this is not freezing, but sitting still/grooming. **F)** Rearing, an exploratory trait in mice where the animal stands on its hind legs with the paws on the walls of the box (PVcre 28.27 ± 6.65, AcanKO 21.1 ± 6.33, p-value = 0.03). PVcre (green) n= 11, AcanKO (purple) n = 10. **Zero maze G)** Percentage of time spent in the open zones of the zero maze. **H)** Total number of entries into the open zones (PVcre 45.30 ± 9.59, AcanKO 30.36 ± 9.31, p-value = 0.003) **I)** Average length (s) of visit per entry into the open zones (PVcre 2.53 s ± 0.73, AcanKO 4.83 s ± 1.51, p-value = 0.005). **J)** Head dips, reaching the head over the edge of the zero maze (PVcre 28.0 ± 5.8, AcanKO 20.73 ± 6.96, p-value = 0.02). **K)** Rearing (PVcre 7.20 ± 3.22, AcanKO 3.45 ± 2.06, p-value = 0.007). Unpaired Student’s t-test. Data shown as mean ± SD. Each dot represents one animal. PVcre (green) n= 10, AcanKO (purple) n = 11.

Our results show that the WFA+ PNN deficient ACANflx/PVcre mice and controls had similar locomotor abilities (Fig. 6A), but the ACANflx/PVcre mice spent more time in the inner zone compared to the PVcre mice (Fig. 6B, PVcre 4.47% ± 1.25, ACANflx/PVcre 8.24% ± 3.05, p-value = 0.002, unpaired Student’s t-test). ACANflx/PVcre mice did not enter the inner zone more often (Fig. 6C), but rather spent more time in the inner zone during each entry (Fig. 6D, PVcre 0.39 s ± 0.17, ACANflx/PVcre 0.68 s ± 0.36, p-value = 0.04). The distance traveled in the inner zone was similar between groups (Fig. S5A), but the speed of the ACANflx/PVcre mice was lower than controls (Fig. S5B, PVcre 0.19 m/s ± 0.07, ACANflx/PVcre 0.13 m/s ± 0.05, p-value = 0.04). We observed that the ACANflx/PVcre mice spent more time in a sedentary state in the inner zone compared to PVcre mice (Fig. 6E, PVcre 0.98 s ± 0.85, ACANflx/PVcre 6.44 s ± 6.99, p-value = 0.02). They did not freeze, but remained stationary and groomed. Additionally, the ACANflx/PVcre mice show less explorative behavior, reflected in the amount of times the mice reared on their hind legs (Fig. 6F, PVcre 28.27 ± 6.65, ACANflx/PVcre 21.1 ± 6.33, p-value = 0.03).

In addition to the open field, the animals’ anxiety levels were tested in the zero maze. The zero maze is a circular track elevated on four legs with four zones: two with walls on both sides, two open. Mice will naturally spend less time in the open zones and more in the closed zones with walls. The groups spent approximately the same amount of time in the open zone (Fig. 6G); however, the ACANflx/PVcre mice had less frequent but longer entries to the open zones (Fig. 6H and I. Entries: PVcre 45.30 ± 9.59, ACANflx/PVcre 30.36 ± 9.31, p-value = 0.003. Time in zone: PVcre 2.53 s ± 0.73, ACANflx/PVcre 4.83 s ± 1.51, p-value = 0.005). Additionally, we found that every time the ACANflx/PVcre mice entered the open zone with just their heads, they would also enter with the rest of their body in 60% of the occurrences; in the control group, this only occurred 34% of the time (Fig. S5C, PVcre 34.68% ± 5.93, ACANflx/PVcre 60.24% ± 20.80, p-value = 0.002). This behavior may reflect a low level of risk assessment among the ACANflx/PVcre mice. Moreover, ACANflx/PVcre mice dipped their heads over the edge of the open zones less frequently, and the occurrence of rearing was also reduced (Fig. 6J and K, head dips: PVcre 28.00 ± 5.85, ACANflx/PVcre 20.73 ± 6.96, p-value = 0.02. Rearing: PVcre 7.20 ± 3.22, ACANflx/PVcre 3.45 ± 2.06, p-value = 0.007). Taken together, the behavior of the ACANflx/PVcre mice in the open field and zero maze indicates that they are less explorative and are less likely to assess the risks in their environment compared to control animals, potentially due to lower levels of anxiety.

### *In vivo* but not *in vitro* chABC treatment of PNN deficient ACANflx/PVcre mice reduces PV+ firing rate, and increases excitatory activity

ACANflx/PVcre mice lack WFA+ PNNs on PV+ interneurons, but intrinsic and firing properties are unaffected (Fig. 3.), in contrast with several papers that find a reduction of firing rate in PV+ cells treated with chABC to digest PNNs (Balmer, 2016; Liu et al., 2022, 2023; Tewari et al., 2018). To investigate if the effects found with chABC are due to other factors than PNN digestion specifically, we treated adult ACANflx/PVcre mice with chABC. We injected chABC (AMSBIO) in V1 three days prior to acute slice preparation, and used V1 in the untreated hemisphere as a control. Surprisingly, PV+ cells in the V1 of the untreated control had a steeper FI curve and a higher maximum firing frequency, than PV+ cells in the chABC treated V1 (Fig. 7E-F). No significant changes were found in intrinsic properties (Fig. 7. A-D), or threshold current (Fig. 7G). Additionally, sEPSC frequency was increased, and sIPSC amplitude was reduced, indicating increased excitatory activity, potentially due to reduced inhibitory activity (Fig. 7H-K).

**Figure 7.**
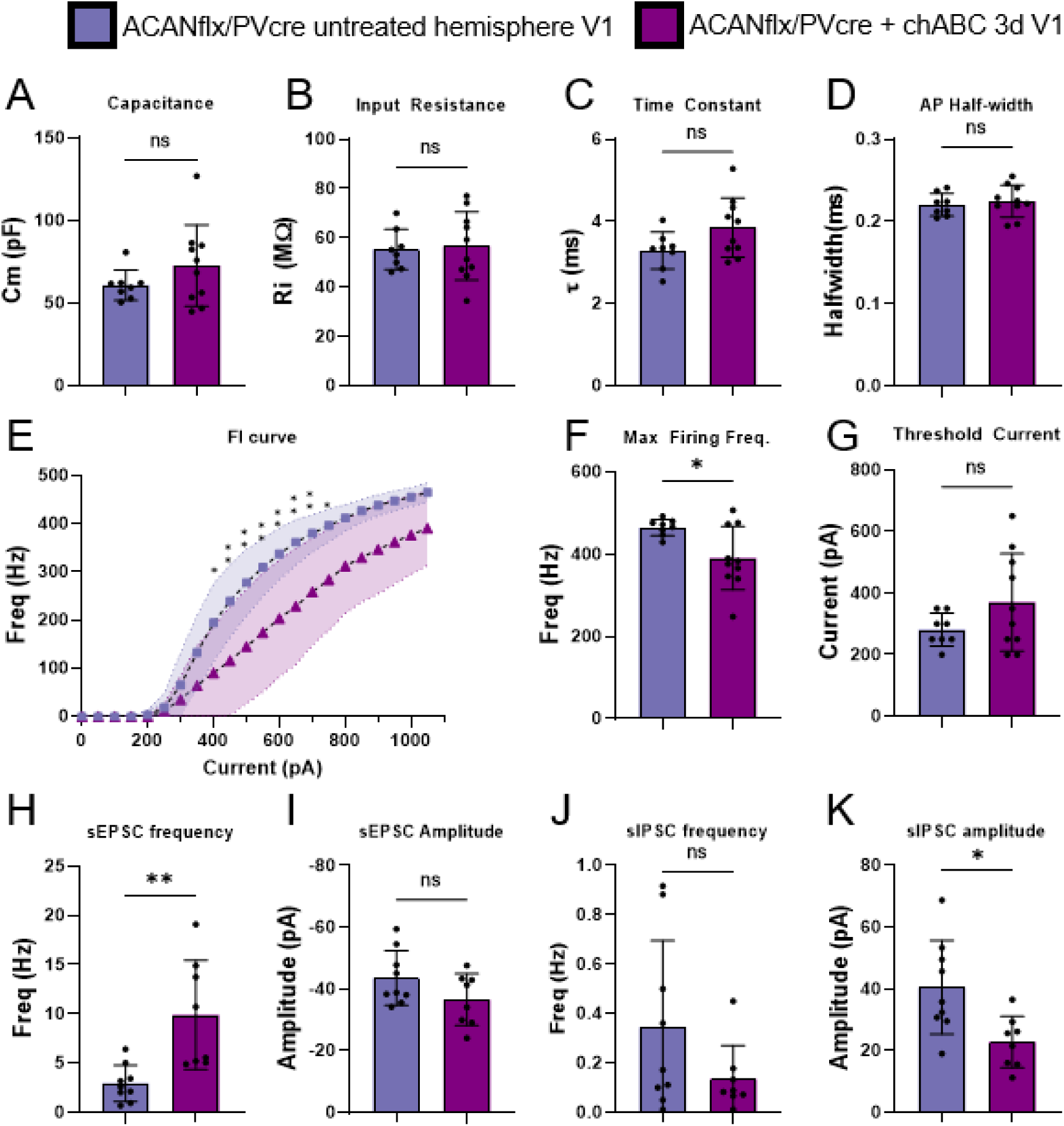
Reduced firing rate and increased sEPSC frequency in V1 PV+ cells in ACANflx/Pvcre mice injected intracerebrally with chABC three days prior to slice preparation, relative to untreated hemisphere. No significant difference in A) capacitance (pF), B) input resistance (MOhm), C) Time constant (Tau, ms) or D) Action potential halfwidth (ms) in chABC treated V1 (purple, n = 10/4 (cells/mice)) vs untreated hemisphere (blue, n = 8/4), student’s t-test (ns = p>0.05). E) FI curve of chABC treated cells rises more slowly and displays a lower firing frequency than in the untreated cells (n = 10/4, 8/4 Sidak’s multiple comparisons, p < 0.05 = *, p < 0.01 = **). F) Maximum firing frequency is also decreased in the chABC treated cells (n = 10/4, 8/4 Welch’s t test, p = 0.0143). G) Threshold current measured in 50pA increments is not changed (n = 10/4, 8/4 Welch’s t test). H) sEPSC frequency is increased in the chABC treated cells (n = 10/4, 8/4 Welch’s t test, p = 0.0089), while I) sEPSC amplitude is unchanged (n = 10/4, 8/4 student’s t-test). J) sIPSC frequency is unchanged (n = 10/4, 8/4 Welch’s t-t est). K) sIPSC amplitude is reduced in the chABC treated cells (n = 10/4, 8/4 student’s t-test, p = 0.101).

We also performed acute chABC treatment during whole cell patch, as in (Tewari et al., 2018). Briefly, PV+ cells were patched and an initial recording of firing properties and sEPSCs and sIPSCs was acquired. Treatment with chABC, or regular recording solution for controls, was applied for 50 minutes before another recording was acquired. These recordings did reveal effects in intrinsic properties, but these effects were also apparent in the ACSF treated controls, and cannot be attributed to chABC treatment (Fig S3).

## Discussion

Development of inhibitory circuits is a fundamental part of the transition from a juvenile to an adult brain state, wherein the assembly of perineuronal nets (PNNs) around PV+ interneurons coincides with the end of maturation. We have studied the outcome of disrupted WFA+ PNN development around PV+ neurons in ACANflx/PVcre mice, a transgenic mouse line with a germline knockout of ACAN in PV+ neurons. While PNNs have been suggested to support PV+ neuron function, e.g. by facilitating their high firing rate (Balmer, 2016; Favuzzi et al., 2017; Lensjø, Lepperød, et al., 2017; Liu et al., 2023; Tewari et al., 2018), we found no differences in the firing frequency of PV+ interneurons in the ACANflx/PVcre line (Fig. 3). It has previously been shown that acute removal of PNNs using chABC in V1 enhances both theta (7-10Hz) and gamma (30-40 Hz) during spontaneous activity (Lensjø, Lepperød, et al., 2017). We did not observe any differences in network activity in V1 (Fig. 4). Additionally, we did not discover any signs of elevated plasticity in the ACANflx/PVcre line regardless of the brain-wide impact on WFA+ PNN development around PV+ neurons. Our contradictory results indicate that previous findings may be caused by the broad effects of chABC, e.g. its digestion of CSPGs in general, and not strictly by the elimination of PNNs. Alternatively, when WFA+ PNNs are knocked out in the germline, compensatory mechanisms during development counteract the effects of the knockout. A similar phenomenon has been shown in a Brevican knockout model, where the germline knockout had limited effects, while shRNA mediated Brevican silencing at P12 caused strong effects on PV+ neuron physiology, and animal behavior (Favuzzi et al., 2017). To circumvent the developmental effect of a germline ACAN knockout, we performed additional experiments on ACANflx mice, not crossed with PVcre, where a PHP.eB PV:cre AAV was injected retro-orbitally to induce an adult, PV+ ACAN knockout. Here, an ocular dominance shift was evident after a brief (four day) monocular deprivation, indicating that the manipulation had generated an elevated level of plasticity in the visual cortex. We did not detect any changes in the local field potential in the adult PV+ ACAN knockout animals.

The PNNs are thought to have a dual role in memory processing. They have restrictive properties that negatively affect memory formation (Blacktop et al., 2017; Carulli et al., 2020; Slaker et al., 2015), while on the other hand, the same properties provide support and stability to the memory consolidation and storage process (Happel et al., 2014; Rowlands et al., 2018; Thompson et al., 2018). Even with a brain-wide depletion of WFA+ PNNs, ACANflx/PVcre mice demonstrated similar memory retrieval performance as controls during both recent (24 hours) and remote (three weeks) memory tests in the Morris water maze task. They did display a slight superiority during training, but behavior analysis indicated that this was due to a low level of anxiety-related behavior that aided them in the training process. It is well documented that anxiety level influences performance in the water maze (Darcet et al., 2014; Higaki et al., 2018; Pritchett et al., 2016). The ACANflx/PVcre mice exhibited comparable performance to controls during reversal training, a process that is unaffected by anxiety because of the animals’ familiarity with the environment.

The difference in anxiety-related behavior was confirmed in the open field and the zero maze. The ACANflx/PVcre also exhibited reduced risk assessment and reduced exploration in the open field and elevated plus maze. This impact on behavior may reflect an excitatory/inhibitory imbalance during development (Sohal et al., 2009); however, we do not have any longitudinal studies on the ACANflx/PVcre mice to confirm this developmental effect. The stark opposite effect was discovered in a recent study where they injected chABC in the medial prefrontal cortex (Li et al., 2024). However, as seen in many other studies, the method of PNN perturbation is influential and makes the results difficult to compare.

As the brain matures from juvenile to adult with a corresponding decrease in plasticity, inhibitory activity increases together with increased PV expression and PNN density (Baker et al., 2017; Caballero et al., 2014). We did not observe any abnormalities in the electrophysiological properties of PV+ neurons or in the recorded circuit activity in the ACANflx/PVcre. In contrast to these findings, we and others have previously shown that enzymatic PNN removal causes changes in LFP power, and strong effects on plasticity (Lensjø, Lepperød, et al., 2017; Pizzorusso et al., 2002). Moreover, we were not able to induce an ocular dominance shift in the ACANflx/PVcre line, while this did occur in the animals where ACAN had been knocked out in adults. Together with the work on the Brevican knockout from Favuzzi et al. (2017), and the strong effects on neuroplasticity in the visual cortex induced by local aggrecan knockout (Rowlands et al., 2018), and PV specific knockout in adult mice, our data suggests that compensatory mechanisms have been initiated to counteract the loss of PNNs in the ACANflx/PVcre line. This may occur as changes in expression of other PNN components or proteases that degrade neural ECM, or changes on the cellular and synaptic level, and should be investigated further. Here, we detected an increase in neurocan and tenascin-r, and a reduction Semaphorin-3a gene expression in the germline KO, that was not found in the adult, virus mediated KO. This suggests that compensation could be mediated by altered expression of other PNN components. Furthermore, in areas where PV+ neuron and PNN density is lower, such as postrhinal cortex, somatostatin expressing interneurons appear to have many of the same functions as PV+ neurons, and show significant similarities in physiological properties to PV+ cells that they do not otherwise have (Sugden & Connors, unpublished observations).

It should also be considered that while germline PV+ ACAN KO ablates WFA and ACAN staining, qPCR analysis on bulk tissue indicates that aggrecan expression is still largely present. This baseline expression of aggrecan from other cell types, paired with increased expression of other PNN components, might produce a sufficient PNN-like structure to rescue effects of ACAN perturbation in PV+ cells. Indeed, when chABC is applied to adult mice lacking ACAN in PV+ cells, PV+ neuron excitability is increased, a phenotype associated with chABC treatment of WT animals. This could further indicate that the compensatory mechanisms are CSPG related, as the compensation is wiped out by chABC treatment, but this remains speculation. The chABC effect on PV+ ACAN KO mice could be interpreted in several ways; maybe complete digestion of all CS in PNNs are required for loss-of-function, or perhaps complete digestion of all CS, not only in PNNs on PV cells are required for strong effects on cells and the network. Regardless, we urge that the wording used to describe chABC treatment in PNN research is adjusted, as it cannot be claimed to be specific to PNNs. Indeed, a seminal paper on PNNs used more restrained language in its abstract: «CSPG degradation with chondroitinase-ABC» (Pizzorusso et al., 2002).

In conclusion, our results showed that germline knockout of ACAN in PV+ neurons is an effective way of disrupting PNN development around this subset of neurons. Yet, the absence of WFA+ PNNs had little to no impact on memory processing or the intrinsic physiological properties of PV+ interneurons in V1. Our discoveries could put into question the role of PNNs in memory processing, and challenge the notion that PV+ neurons are reliant on PNNs as a facilitator. However, the differences between the ACANflx/PVcre animals and adult animals where ACAN had been acutely knocked out in PV+ cells. This suggests that compensatory mechanisms may be initiated to sustain normal function in the absence of PNNs, and calls for further research on genetic PNN perturbation in the germline and in adults, as well as the true effects of chABC treatment.

## Acknowledgements

We thank Tove Klungervik at the Department of Bioscience for assistance with genotyping and to Hong Qu at the Institute of Basic Medical Sciences for assistance with slide scanning.

This work was funded by the Research Council of Norway (project #250259 and 549217 to MF and 685248 to TH,) the Norwegian Health Association (16036 to EHT) and the University of Oslo’s Strategic Research Initiative CINPLA.

## Conflict of interest

The authors declare no conflict of interest.

## Author contributions

SG and EHT contributed equally as first authors. SG, EHT, TH, MF and KKL designed the study, SG and TM performed in *vitro* electrophysiology experiments and analysis, MBR performed in *vivo* electrophysiology experiments after pilots were performed by KKL, EHT performed behavioral testing and immunohistochemistry, SG and GHV performed cloning and virus preparation, GHV and IN performed qPCR, IN performed RO injections and tissue extraction for qPCR, MBR, AB, EHT and TH performed analysis of in *vivo* electrophysiology. SG, EHT and KKL wrote the first draft of the manuscript, with input from all authors. All authors have seen and approved the final version of the manuscript.

## Material and Methods

### Ethics and approvals

All experimental procedures were conducted in agreement with guidelines for work with laboratory animals described by the European Union (directive 2010/63/EU) and the Norwegian Animal Welfare Act of 1974/2010 (replaced by the new Animal Welfare Act in 2010). Experiments were approved by the National Animal Research Authority of Norway (Mattilsynet). Experiments followed the principles of humane experimental techniques, to Replace, Reduce and Refine.

### Animals

The ACANflx/PVcre mice were obtained by crossing homozygous ACAN-flox (ACANflox^+/+^) (Rowlands & Lensjø et al., 2018) mice with homozygous PVcre mice (PVcre^+/+^) (Hippenmeyer et al., 2005). From this first round of breeding, all mice had an ACANflx^+/-^/PVcre^+/-^ genotype and went through another cross to obtain the homozygous genotype. Only one copy of the Cre allele is necessary for recombination. Therefore, from the second-generation litters, we used both ACANflx^+/+^/PVcre^+/+^ and ACANflx^+/+^/PVcre^+/-^. Animals were bred locally in the animal facility at Dept. of Comparative Medicine at the University of Oslo. PVcre mice (+/- and +/+) were used as controls. ACANflx^+/+^ were included in the experiments on visual cortex plasticity. The animals were between three and six months at the time of experiments and housed in individually ventilated cages (IVC, GM500, Scanbur) with a 12h light/dark cycle. Behavioral experiments were conducted in the dark phase.

### Extracellular recording using silicon probes

Surgery was performed under isoflurane anesthesia (5 % induction, 1-2 % maintenance) using a SomnoSuite vaporizer (Kent Scientific). The animal was given a local subcutaneous injection of Bupivacaine adrenalin (Marcain 0.5%, 1 mg/kg) in the scalp and a subcutaneous injection of Buprenorphine (Temgesic, 0.05 mg/kg) before the start of the surgery. The animal was secured in a stereotaxic surgical frame using ear bars placed anterior to the ear canal. The eyes were covered with Viscotears (2mg/g, Vitus Apotek) to prevent dehydration. A small patch of skin was removed to expose the skull, and the skull was lightly scored using a surgical blade. A small hole was drilled into the skull using a hand-held Perfecta-300 dental drill (W & H Nordic) to fit in a screw used as a ground reference during the electrophysiological recording. A custom head-bar was glued to the skull using Vetbond (3M, VWR), and secured using C&B SuperBond dental acrylic (Parkell). The animal was given a subcutaneous injection of Norodyl (carprofen, 5mg/kg) and placed in a cage to recover. Norodyl was given for three consecutive days after surgery.

Animals were habituated to the running wheel and trained to run voluntarily for two days prior to the extracellular recording experiment. On the day of the first recording session, a large craniotomy was made over the primary visual cortex (V1) centered at 3.1 mm laterally from lambda. Following the craniotomy, the exposed brain was covered with gel foam soaked in sterile saline solution, the animal was transferred to the experimental setup and allowed to recover from the brief anesthesia. A silicone probe (NeuroNexus A1x32-5mm-25-177) was attached to a stereotaxic frame and slowly lowered ∼800-900 µm into V1. Before the recordings started, the probe was left in the tissue for a minimum of 30 minutes to let the tissue recover (Niell & Stryker, 2008).

Animals were tested for ocular dominance plasticity by using monocular deprivation (MD). Immediately after the first round of extracellular recordings, or if the animals were only used for “after MD” recordings, a drop of Alcaine was administered to the contralateral eye before it was sutured close. After 4 days, extracellular activity was recorded shortly after removing eyelid sutures. If it was the last recording session, the probe was coated with a fluorescent dye (Sigma-Aldrich 42364) for post-mortem histological verification, and the animal was transcardially perfused as described below.

The 32-channel silicone probe was connected to an Intan RHD 2132 amplifier board (hardware bandpass filtering between 1.1 Hz and 7.5 kHz) which was connected to an Open Ephys acquisition system (Siegle et al., 2017) via a thin SPI cable (Intan Technologies, USA). Data was sampled at 30 kHz and saved to disk for offline analysis. We used Expipe (Lepperød et al., 2020) to manage all data which were then saved to Exdir format (Dragly et al., 2018) for further processing. All data were low-pass filtered at 300 Hz, processed with a notch filter to remove 50 Hz line noise (quality factor Q=100, scipy.signal.iirnotch function), resampled at 1 kHz, and z-scored according to channel. Visual stimulus periods were extracted and aligned according to stimulus onset for the LFP trace and frequency spectrum. To investigate the LFP oscillations for the different frequency bands during spontaneous and stimulus evoked LFP activity, we computed the spectrogram of the z-scored LFP signals with continuous wavelet transform applied using a Morlet wavelet (Lee & Choi, 2019). The median power spectral density was then extracted for 200 ms before and 200 ms after stimulus onset.

### Visual stimulation

Visual stimuli was generated using custom scripts written in Psychopy (Peirce, 2007) and presented on a computer monitor placed 25 cm in front of the mouse, spanning 30° degrees of the horizontal visual field. Stimuli consisted of full-field drifting sinusoidal gratings of 100 % contrast, with a temporal frequency of 2 Hz and spatial frequency of 0.04 cycles/degree. The gratings were presented at eight orientations (separated by 45°) for 2 s each, followed by 1 s of mean-luminance gray. Trials were randomized, and eight repetitions of each orientation were used per electrophysiological recording with gratings.

### AAV construct cloning and packaging

For patch-clamp electrophysiology experiments, mice (p35-70) were injected intravenously, in the retro-orbital sinus, with a systemic PHP.eB AAV expressing flex-tdTomato, to label PV+ cells. For adult KO of ACAN in PV+ cells, ACANflx mice were injected retro-orbitally with a PV enhancer Cre-P2A-mGreenLantern or GFP virus. The AAVs were produced as described in a protocol by Challis et al. (Challis et al., 2019). Briefly, AAV HEK293T cells (Agilent) were cultured in DMEM with 4.5 g/L glucose & L-Glutamine (Lonza), 10% FBS (Sigma) and 1% PenStrep (Sigma), in a 37 °C humidified incubator. The cells were thawed fresh and split at ∼80% confluence until four 182.5 cm2 flasks were obtained for each viral prep. The cells were transfected at 80% confluence and the media was exchanged for fresh media directly before transfection. The cells were triple transfected with the desired construct, dF6 helper plasmid and PHP.eB serotype plasmid. Polyethylenimine (PEI), linear, molecular weight (MW) 25,000 (Polysciences, cat. no. 23966-1) was used as the transfection reagent. Media was harvested three days after transfection and kept at 4 °C, and media with cells was harvested five days after transfection and combined with the first media harvest. After 30 min centrifugation at 4000 × g, the cell pellet was incubated with SAN enzyme (Arctic enzymes) for 1 h. The supernatant was mixed 1:5 with PEG and incubated for 2 h on ice, then centrifuged at 4000 × g for 30 min to obtain a PEG pellet containing the virus. The PEG pellet was dissolved in SAN buffer and combined with the SAN cell pellet for incubation at 37 °C for 30 min. To purify the AAV particles, the suspension was centrifuged at 3000 × g for 15 min. The supernatant was loaded on the top layer of an Optiseal tube with gradients consisting of 15%, 25%, 40% and 60% iodixanol (Optiprep). Ultracentrifugation was performed for 2.5 h at 18 °C at 350,000 × g in a type 70 Ti rotor (Beckman Coulter). The interface between the 60 and 40% gradient was extracted along with the 40% layer, avoiding the protein layer on top of the 40% layer. The viral solution was filtered through a Millex-33 mm PES filter and transferred to an Amicon Ultra-15 centrifugal filter device (100-kDa molecular weight cutoff, Millipore) for buffer exchange. A total of four washes with 13 ml DPBS were performed at 3000 × g before concentration to a volume of ∼500–750 µL. Viral solutions were sterilized using a 13 mm PES syringe filter 0.2 μm (Nalgene), and stored in sterile, low-bind screw-cap vials at 4 °C.

Viral titers were determined using qPCR with primers targeting AAV2 ITR sites. Briefly, 5 µL of viral sample was added to 39 µL ultrapure H2O, 5 µL 10× DNase buffer, 1 µL DNase, and incubated at 37 °C for 30 min to eliminate all DNA not packaged into AAV capsids. 5 µL of the DNase-treated sample was added to a reaction mix consisting of 10 µL 2× SYBR master mix, 0.15 µL of each primer (100 µM) and 4.7 µL nuclease free H2O. Cycling conditions for the qPCR program were: 98 °C 3 min/98 °C 15 s/58 °C 30 s/read plate/ repeat 39× from step 3/melt curve.

The PHP.eB serotype plasmid, pUCmini-iCAP-PHP.eB was a gift from Viviana Gradinaru (http://n2t.net/addgene:103005;RRID:Addgene_103005). The DeltaF6 helper plasmid, pAdDeltaF6 was a gift from James M. Wilson (http://n2t.net/addgene:112867;RRID:Addgene_112867). pAAV-FLEX-tdTomato was a gift from Edward Boyden (Addgene plasmid # 28306, RRID:Addgene_28306). PV:mGreenLantern-P2A-NLS-CRE was cloned via multiple intermediate plasmids, is available upon request, and will be deposited at Addgene.

AAVs were injected intravenously in the retro-orbital sinus. Briefly, each mouse was anesthetized in 5% isoflurane and injected retro-orbitally with 5×10^12^ vector genomes of AAV. A drop of Alcaine was administered to the eye post-injection, and the mice were left to recover from anesthesia in a heated chamber for a few minutes.

### Acute slice preparation

Acute slices were prepared from mice aged 70-140 days according to the protective recovery method described by Ting et al. (Ting et al., 2018). Mice were anesthetized in 5% isoflurane and injected intraperitoneally with a lethal dose of pentobarbital. When breathing had slowed and the animal no longer responded to strong toe or tail pinching the mouse was transcardially perfused with freshly bubbled, chilled NMDG-ACSF, containing the following in mM: 93 NMDG, ∼93HCl, 2.5 KCl, 1.2 NaH_2_PO_4_, 30 NaHCO_3_, 20 HEPES, 25 glucose, 5 sodium ascorbate, 2 thiourea, 3 sodium pyruvate, 10 MgSO_4_.7H_2_O, 0.5 CaCl_2_.2H_2_O. The brain was dissected and mounted on a Leica vt1200s vibratome in a continuously cooled and bubbled chamber. Coronal sections (300 µm) were cut in a caudal to rostral direction until V1 was no longer visible. Slices were left to recover for 25 minutes in heated NMDG-ACSF while sodium was gradually reintroduced. Following recovery slices were moved to a high HEPES holding ACSF containing the following in mM: 92 NaCl, 2.5 KCl, 1.2 NaH_2_PO_4_, 30 NaHCO_3_, 20 HEPES, 25 glucose, 5 sodium ascorbate, 2 thiourea, 3 sodium pyruvate, 2 MgSO_4_.7H_2_O, 2 CaCl_2_.2H_2_O, at room temperature. The slices were incubated in the recovery chamber for at least one hour before transfer to the recording chamber. All solutions were continuously bubbled with 95% O_2_, 5% CO_2_.

### Patch-clamp

Whole-cell patch-clamp recordings were performed in V1. Cells were visualized on a Zeiss axioscope using infrared DIC illumination and a hamamatsu c2400 CCD camera. The PV+ cells were identified by the virally expressed, red fluorescent tdTomato. The recording chamber was continuously perfused at a rate of 4ml/min with continuously carbogen-bubbled recording ACSF of the following composition in mM: 124 NaCl, 2.5 KCl, 1.2 NaH_2_PO_4_, 24 NaHCO_3_, 5 HEPES, 12.5 Glucose, 2 MgSO_4_.7H_2_0, 2 CaCl_2_.2H_2_0. pH was adjusted to 7.3-7.4, and osmolarity to 305-310 mOsm. Pipettes were pulled from thick-walled borosilicate glass capillary tubes to a tip resistance of 3.5-4.9 MΩ. For current clamp experiments in Fig 3, pipettes were filled with an intracellular solution containing the following, in mM: 120 K-Gluconate, 20 KCl, 10 EGTA, 10 HEPES, 2 MgCl_2_, 2 Na_2_ATP (Hu et al., 2014). pH was adjusted to 7.3 using KOH, and osmolarity to ∼295 mOsm. For current and voltage clamp experiments in Figure 7, S2 and S3, a low-chloride IC solution was used for simultaneous measurement of sEPSCs and sIPSCs: 140 K-gluconate, 2 KCl, 5 EGTA, 10 HEPES, 4 Mg-ATP, 0.5 Na-GTP, 5 creatine phosphate (Tsintsadze et al., 2017). Signal was amplified and low-pass filtered on a Multiclamp 700b amplifier (Molecular devices; 6kHz), and digitized on a Digidata 1550B digitizer at 100kHz (Molecular devices) using the pClamp11 software package (Molecular Devices).

Membrane potential was measured immediately upon establishment of the whole-cell configuration in I=0 mode. Cells with access resistance (Ra) greater than 25 MΩ were discarded. Maximum firing rate and FI curves were established from a 1s step current stimulus, starting at -100pA and increasing by 50pA per sweep. The start of each sweep featured a 500ms hyperpolarizing stimulus, from which the input resistance (Ri), time constant (*τ*) and capacitance (Cm) was measured.

Current clamp recordings were analyzed using the pyABF module (Harden, 2022) for ABF file handling and visualization, and the eFEL module (Geit et al., 2018) for feature detection in custom python 3.6 scripts (available upon request). Input resistance was calculated from the voltage deflection of a hyperpolarizing current step. The membrane time constant (*τ*) was calculated from an exponential decay fit to the same hyperpolarizing current step. Capacitance (Cm) was calculated from the slowest time constant and the input resistance (Golowasch et al., 2009). For establishing the FI curve and the maximum firing rate, eFEL was used to count action potentials during the 1s stimulus using a derivative threshold of 15 and an interpolation step of 0.01. Action potential detection was plotted and visually verified for at least the fastest spiking trace in each experiment. sEPSCs and sIPSCs were detected and analyzed from 300s (Fig 7.) or 75s (Fig S3) gap free recordings using template matching in Easy Electrophysiology, and custom python scripts (available upon request).

For in vivo chABC treatment of ACANflx/Pvcre mice (Fig 7.), surgery was performed as described above, three days prior to acute slice preparation. 500 nanoliters of 60U/ml chABC (AMSBIO) was injected unilaterally in V1 using a Nanoject III (Drummond). The opposite hemisphere served as an untreated control.

For acute chABC treatment (Fig S2, S3), initial recordings (time 0) were acquired 15 minutes after whole cell configuration was achieved. chABC (Sigma or AMSBIO) was dissolved in 10ml of freshly bubbled ACSF, for a concentration of 1U/ml. Following CC and VC recording at time 0, the recording solution was replaced with continuously bubbled chABC ACSF. Additional recordings were acquired at various time points, including 50 minutes.

### Behavior and learning assessment tests

#### Morris water maze

The pool used was 120 cm in diameter, with 30 cm from the edge of the pool to the platform and with a platform diameter of 11 cm. For analysis and training purposes, the pool was divided into four equal quadrants. The pool was filled with water holding a temperature 13 degrees below body temperature (∼23 degrees). The water was made opaque using water-based white paint (Ready mix Tempera, Panduro). During training the platform was 2 cm under the water surface.

The mice were trained for seven consecutive days with four trials per day. Each mouse was placed in the water close to the wall of the pool in a different quadrant each trial. The order of which quadrant the mice were placed in was changed every day. The mice were allowed to swim freely for 60 seconds before being led to the platform and forced to sit on the platform for around 5 seconds before being taken out. One trial was completed for all the animals before starting the next trial, to ensure an intertrial interval of around 15 minutes for each mouse.

The first probe trial, without the platform, was performed 24 hours after the last training trial. The mice were placed in the pool for 30 seconds before being taken out. A second probe trial, with the same procedure as above, was performed 21 days after the last training day. Training with a new platform location (reversal training) was started after the 21-day probe trial. Reversal training was conducted by repeating the same protocol, but with the platform in a new position in a different quadrant. The animal’s movements in the pool were tracked by Bonsai and analyzed using MATLAB.

The swim patterns of the mice were examined and classified according to six different swimming strategies (Cooke et al., 2019; Illouz et al., 2016). The swim strategies are given a cognitive score. The strategies and their scores: thigmotaxis (1), random search (2), scanning (3), indirect search (4), direct search (5), direct path (6). Based on these strategies, cognitive scores were estimated for each day. The cognitive score is the sum of scores in the four trials divided by four to get the mean. Mice that did not swim, but rather floated in the pool for the duration of the trial, were excluded from the analyses.

#### Open field

The open field box measured 50×50×50 cm. On the day, the animal’s home cage was placed in the room together with the open field box for one hour before testing. The maze was cleaned with ethanol between each session. The animal was placed in the middle of the box, and could explore it freely for five minutes while its movements were monitored using the tracking software ANY-maze. For analysis purposes, the box was divided into the outer- and the inner zone, the inner zone was further divided into the center zone and the middle zone. The inner zone was an approximately 30cm × 30cm large square, the center zone a 10cm × 10cm square inside the inner zone. However, most of the analysis was done comparing the inner zone as a whole with the outer zone. Each mouse was only tested one time.

#### Zero maze

The zero maze is a circular track elevated on four legs. It is divided into four zones, two are open, and the other two have walls on the side of the track. Rodents will naturally seek darker areas and will therefore spend most of their time in the maze in the zones with walls. On the day of testing, the cage was placed in the room with the zero maze for approximately an hour before testing. The mouse being tested was placed in one of the walled zones and was then left to explore the maze for five minutes. Movements were monitored using the tracking software ANY-maze. The maze was cleaned with ethanol between each session so that odor from one mouse would not create a distraction for the following mouse. The mice were only put in the maze once, to inhibit habituation.

### Bulk tissue qPCR

Bulk tissue was extracted from V1 with a tissue puncher, put in RNAlater solution (Invitrogen) and stored at 4℃ until RNA extraction. For RNA isolation, the tissue piece was added to 600 ul RLT lysis buffer from the RNeasy kit (Qiagen) with added 1% (v/v) 2-Mercaptoethanol. The tissue was homogenized using a TissueLyser II instrument at full speed for 3 minutes with one stainless steel bead in each tube. The tubes were centrifuged at 4℃ at 20’000x g for 30 min, and the supernatants were transferred to new tubes. RNA was isolated with the RNeasy mini kit (Qiagen). 1000 ng of RNA from each sample was transcribed to cDNA with FIREScript reverse transcriptase (Solis Biodyne) and random hexamers, using a temperature program of 25℃ 10 min, 50℃ 30 min, 85℃ 5 min, 4℃. The cDNA was diluted 1:5 with nuclease free water and stored at -20℃. Two separate rounds of reverse transcription were performed on the RNA, to limit variance in reverse transcription efficiency.

Samples from a total of 22 mice were analyzed; 9 from the ACANflx/PVcre group, 6 from the ACANflx/AAV-PVcre group, and 7 from the PVcre control group. All samples were run on the same plate for each primer specific mix. A volume of 10 ul was used in each well, with 2 ul cDNA, 0.5 ul of each primer (10 uM), and 5 ul LightCycler SYBR Green Mastermix 2X (Roche). LightCycler 96-well plates were run on a LightCycler 96 machine (Roche) with an activation step of 95℃ 120 sec; denaturation 95℃, 10 sec; primer annealing 58℃, 10 sec; extension 72℃, 7 sec. Melt curve analysis was performed to check product specificity, and any values without a clean peak were removed. All samples were run in duplicates or triplicates, averaged, and normalized to the combination of GAPDH and Cyclophilin. Data analysis was performed with the steps described in figure 5 of the publication from Taylor et al., similar to the ΔΔCq method, to acquire a relative change in expression between the biological sample groups. Tukey’s multiple comparisons test was performed with GraphPad Prism 9.1 software.

qPCR primers can be found in Table S1.

### Immunohistochemistry

An overdose of pentobarbital sodium (Euthasol, 100 mg/kg) was administered as an intraperitoneal injection under isoflurane anesthesia. When breathing had slowed and the animal no longer responded to toe or tail pinching, the heart was exposed and perfused transcardially using 10 mL 1X PBS followed by 10 mL of 4 % PFA (Sigma, F8775, diluted in 1X PBS). For the brain tissue used for qPCR and western analysis, only cold PBS was used for perfusion. Tissue from the visual cortex was dissected out using a tissue puncher and put on a RNAlater solution (Invitrogen). The brain intended for immunohistochemistry was dissected out and left in 4% PFA for approximately 24 hours, before it was transferred to a 30% sucrose solution (prepared in 1X PBS) and left for two days at four ℃. Sectioning was done using a cryostat (Leica, CM1950), creating 40 µm sections that were transferred to 1x PBS with 0.02% sodium azide for later use in immunohistochemistry. To stain for Hyaluronic acid (HA), Neurocan, Brevican, Tenascin-R and Aggrecan, we initially performed antigen retrieval to un-mask antigens after aldehyde fixation. The sections were placed in a tube with 10 mM sodium citrate solution (pH 8.5-9.0) and left in a water bath maintained at 50 Celsius for approximately 45 minutes (Jiao et al., 1999). By using the water bath, free floating sections may be heated to a set temperature ensuring minimal damage. The sections were then left in the citrate buffer solution until it reached room temperature. Sections were washed for 3×5 min in 1x PBS on a shaker. Then they were incubated in a blocking solution (3% BSA, 1.5% Triton X-100, in PBS) for one hour. Sections were transferred to the primary antibody solution (1:500 WFA (Sigma-Aldrich, #L-1516)/1:1000 rabbit/goat anti-PV (Swant, #PV27, #PVG-214)/ rabbit anti-ACAN (Merck Millipore, #ab1031), in block) for 24 hours at room temperature or 48h at 4℃. On day two, sections were washed for 3×5 minutes with 1x PBS before transferring to a secondary antibody solution (streptavidin (#S-11223)/donkey anti-rabbit (#ab150075)/donkey anti-goat (#A-21447), Life Technologies, all with 1:1000) incubated for ∼3 hours. Sections were washed again for 3×5 min with 1x PBS, mounted on Superfrost glass slides, let dry, coverslip using FluorSave™ (Merck Millipore, #345789), and left to cure overnight before imaging.

### Imaging

A confocal spinning-disc microscope (Dragonfly, Andor technologies, Oxford Instruments Company) was used to acquire Z-stacks, using Fusion software (Bitplane). The objective (Nikon) used was 20x, dry, NA = 0.75. Each stack was 613 x 613 µm through a volume of 20 µm, made up of individual images acquired with ∼0.2 µm spacing in the z direction. Images were taken from the medial prefrontal cortex (AP ∼1.7, DV ∼2.3, ML ∼0.3), retrosplenial cortex (AP ∼-1.94, DV ∼1.0, ML ∼0.3-0.7), visual cortex (AP ∼-3.0, DV ∼0.8, ML ∼3.0) (coordinates relative to bregma). Tile images were taken using the AxioScan Z1 slide scanner with the ZEN lite Blue software (Carl Zeiss). Z-stacks were flattened by creating a maximum intensity projection in ImageJ. Cellpose was used to automatically segment PV+neurons for analyzing numbers of cells (Stringer et al., 2021). ImageJ was used for the manual counting of overlap between WFA+ and PV+ neurons. The counting was performed by a blinded subject.

### Statistics and graphical design

Statistics and figures were made using GraphPad Prism 8 or 9. Statistical significance was assessed using either a repeated measures ANOVA/Mixed effect analysis with a Sidak’s multiple comparisons test, an unpaired student t-test of variance or a Kolmogorov-Smirnov test for the FI curves. Significance level were marked as * p < 0.05, ** p < 0.01, and *** p < 0.0001. Biorender.com was used to create the illustrations.

## Notes

### Competing Interest Statement

The authors have declared no competing interest.

