## Supplementary Figures for "Deciphering the Role of Aggrecan in Parvalbumin Interneurons: Unexpected Outcomes from a Conditional ACAN Knockout That Eliminates WFA+ Perineuronal Nets"

**This document contains supplementary figure S1-S5, and Table S1**

**Figure S1**

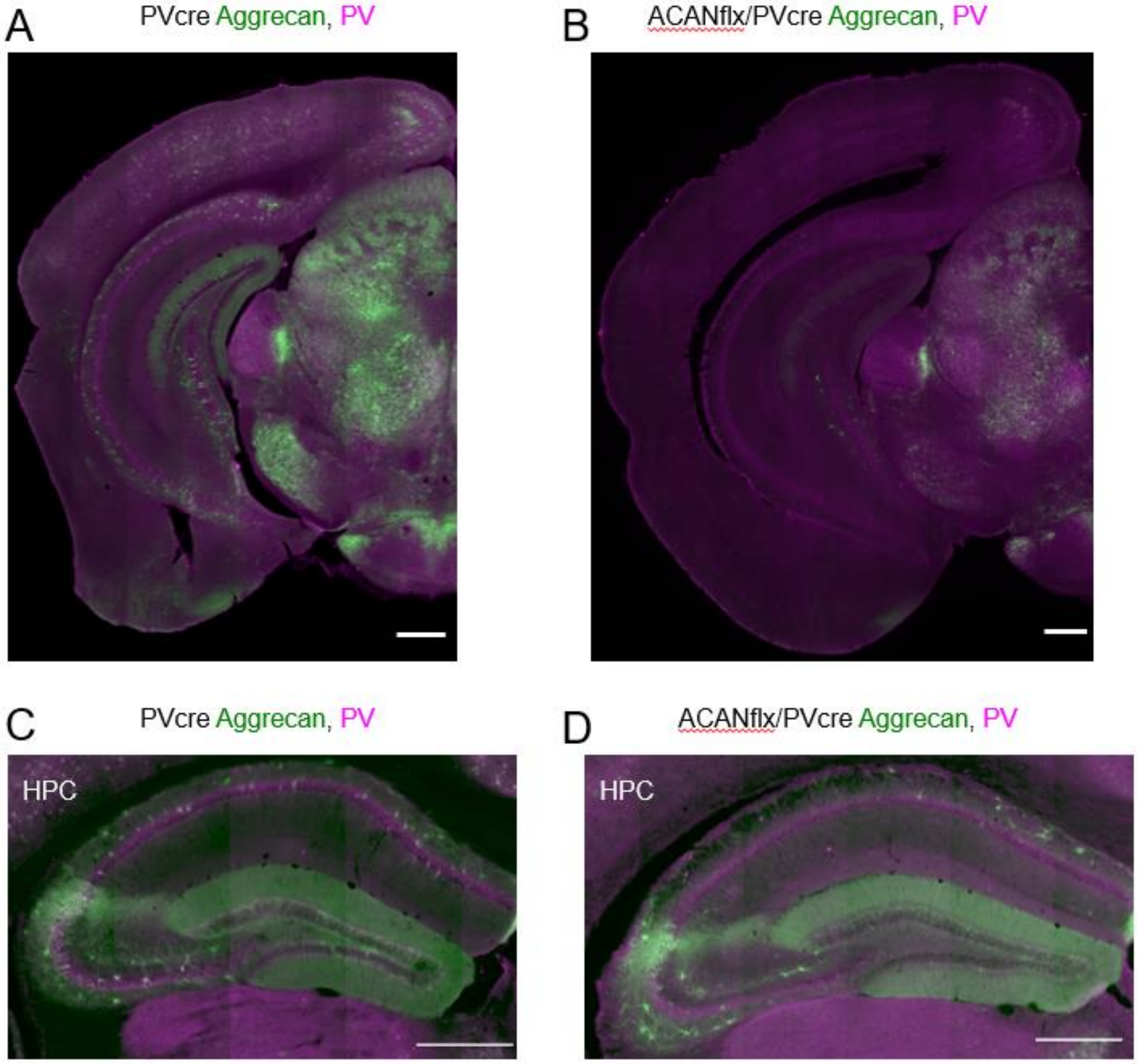


S1. *The Acan knockout (PVcre/Acan-loxP cross) causes a brain-wide depletion of Aggrecan+ PNNs around PV+ neurons. A) Coronal section from a PVcre mouse stained with* Aggrecan *(green) and PV (magenta). B) Coronal section from a ACANflx/Pvcre mouse stained with Aggrecan (green) and PV (magenta). C) Coronal section showing the hippocampus (HPC) of a PVcre mouse stained with Aggrecan (green) and PV (magenta). D) Coronal section showing the hippocampus (HPC) of a ACANflx/PVcre mouse stained with* Aggrecan *(WFA) (green) and PV (magenta). Scale bar = 500 µm*

### Figure S2

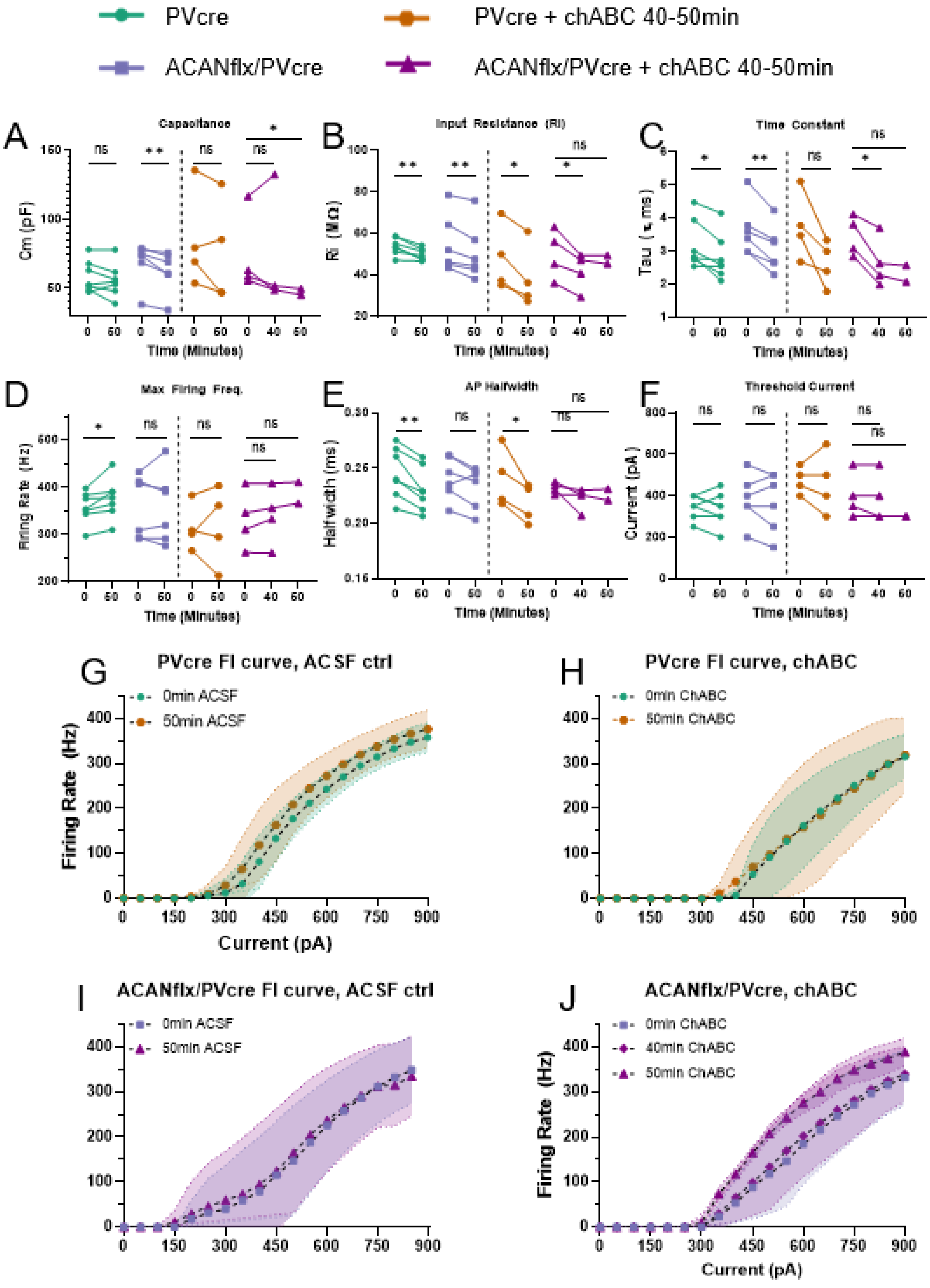


**Figure S2**. **PV+ neurons in acute slices from Pvcre or ACANflx/Pvcre mice treated for 50min with either plain ACSF or ACSF+1U/ml chABC display altered intrinsic electrophysiological and firing properties.** ***A)*** *Capacitance (CM) is significantly reduced in both ACANflox/PVcre treated with ACSF.* ***B)*** *Input resistance (Ri) is significantly reduced in all groups.* ***C)*** *Time constant is reduced in all groups but PVcre + chABC.* ***D)*** *Max Firing Frequency (Hz) is increased in ACSF-treated PVcre, but not significantly changed in any other group.* ***E)*** *Action potential half-width is reduced in PVcre treated with ACSF and chABC.* ***F)*** *Threshold current is unchanged in all groups.* ***G-J)*** *FI curves are unaffected by all treatments. All comparisons were made with paired t-tests. PVcre n = 7 cells, 5 animals, ACANflox/PVcre n = 6 cells, 5 animals, PVcre+chABC n = 4 cells, 4 animals, ACANflox/PVcre-40min n = 2 cells, 2 animals, ACANflox/PVcre-50min 4 cells, 3 animals.*

**Figure S3**

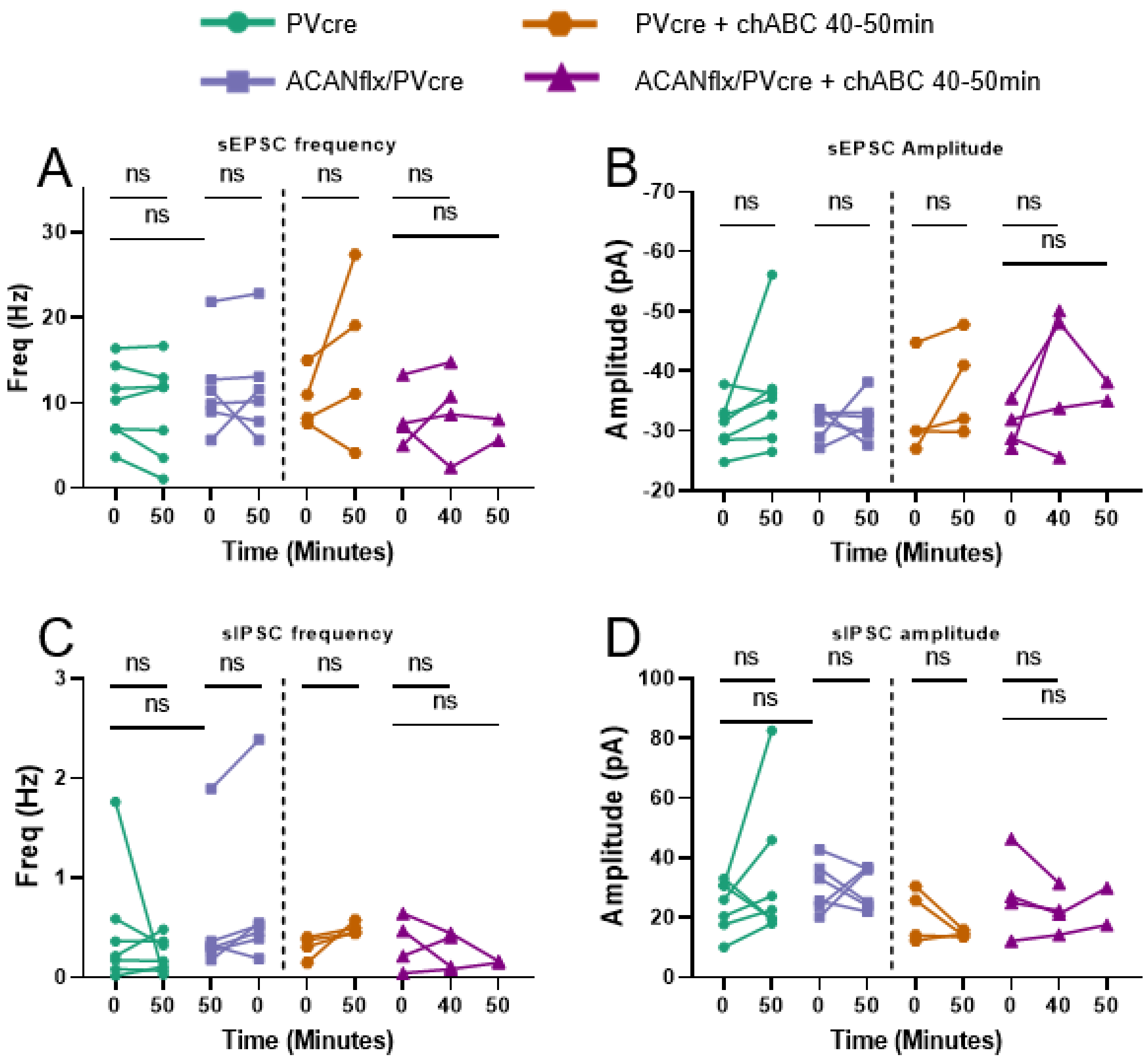


**Figure S3**. **sEPSC and sIPSC frequency and amplitude is unchanged in PV+ cells in acute slices from Pvcre or ACANflx/Pvcre mice treated for 50min with either plain ACSF or ACSF+1U/ml chABC.** ***A-D)*** *No changes were found in sEPSC or sIPSC frequency or amplitude in any treatment groups. All comparisons made with paired t-tests. PVcre n = 7 cells, 5 animals, ACANflox/PVcre n = 6 cells, 5 animals, PVcre+chABC n = 4 cells, 4 animals, ACANflox/PVcre-40min n = 2 cells, 2 animals, ACANflox/PVcre-50min 4 cells, 3 animals.*

**Figure S4**

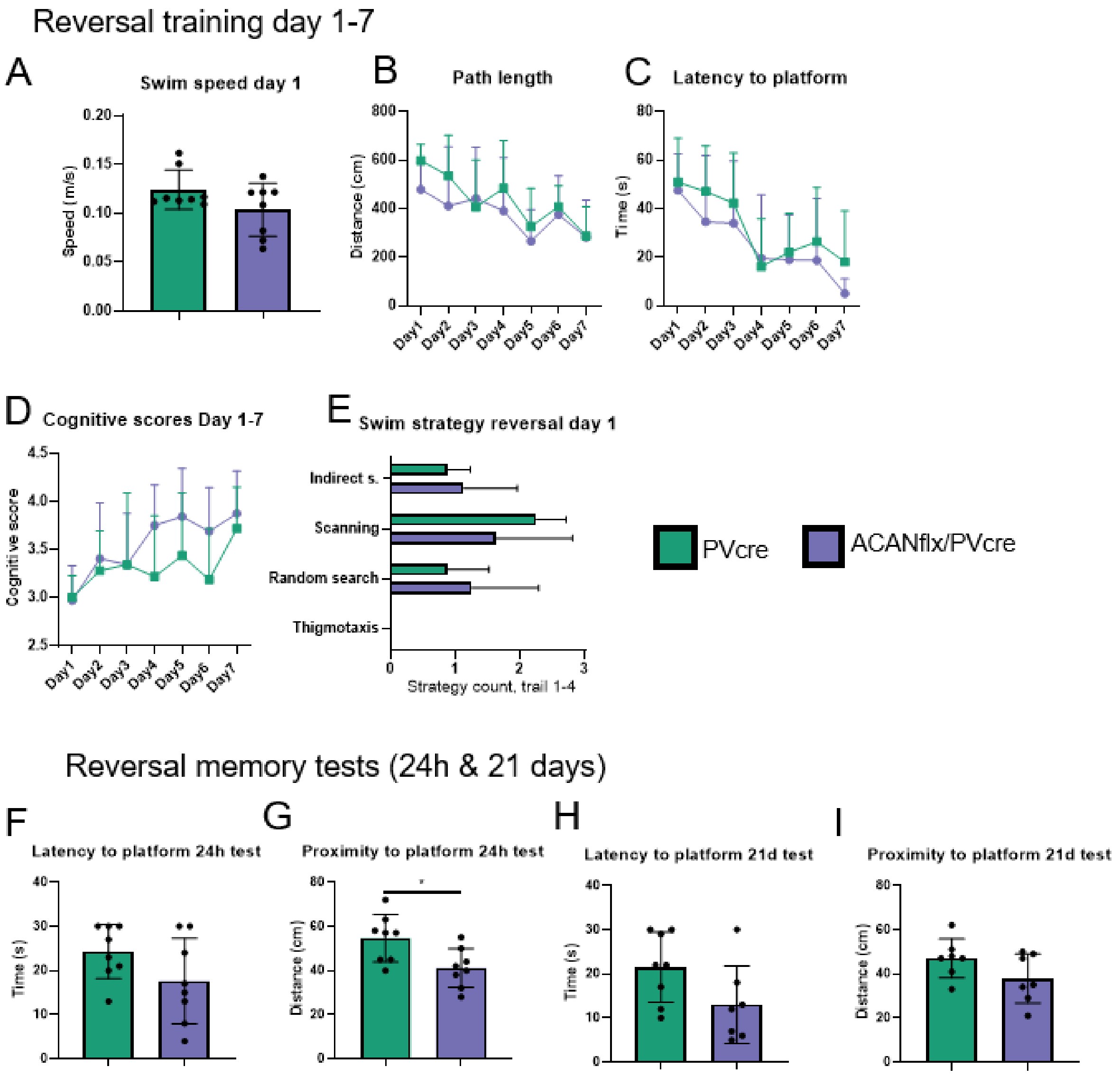


**Figure S3 Reversal training and testing in Morris water maze** *A) Swim speed on day one. B) Path length (cm) during reversal training day 1-7 (Row factor (days of training), p-value = 0.001). C) Latency to platform (s) during reversal training day 1-7 (Row factor (days of training), p-value < 0.0001). D) Cognitive scores given from day 1-7 of reversal training. E) Swim strategy day 1 of reversal training. No occurrences of thigmotaxis. F) Latency (s) to platform during 24 hour memory test G) Proximity (cm) to platform during 24 hour test (mean proximity (cm) PVcre 54.63 cm ± 9.9, ACANflox/PVcre 41.13 cm ± 8.3, p-value = 0.02). H) Latency (s) to platform during the 21 day test. I) Proximity (cm) to platform. Repeated measures ANOVA with Sidak's multiple comparisons test used for A, B and D. Unpaired Student's t-test used for C and F-I. Data shown as mean ± SD. Each dot represents one animal. PVcre (green) n= 8, ACANflox/PVcre (purple) n= 8.*

**Figure S5**

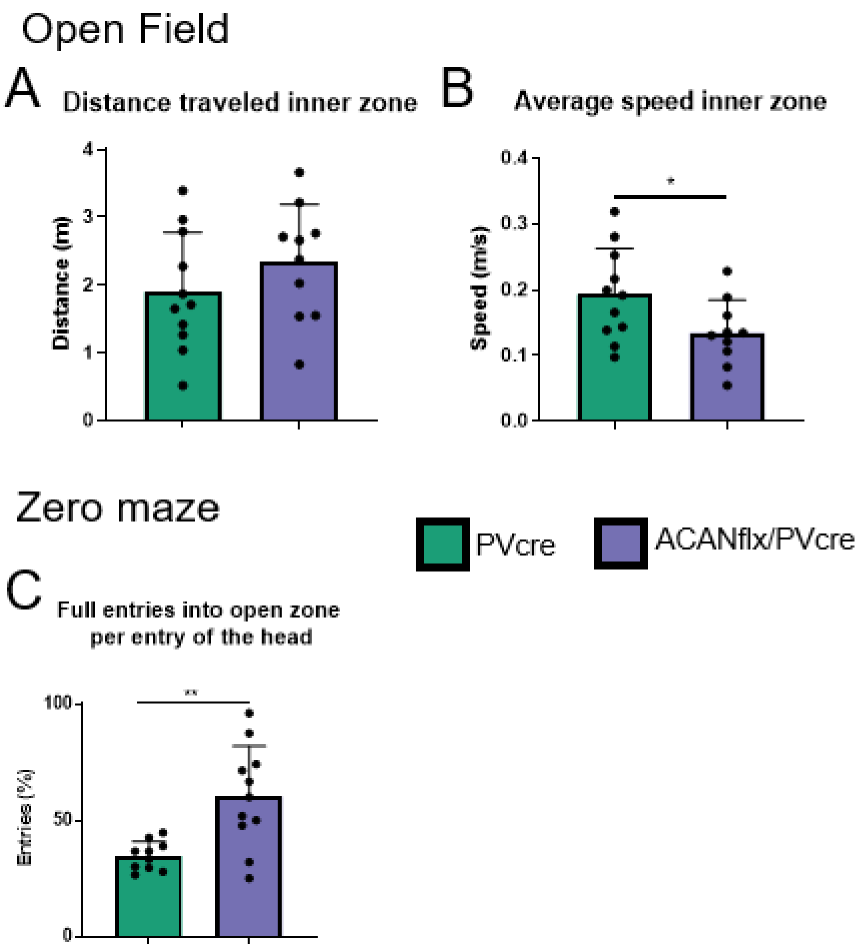


**Figure S4.** ***AcanKO demonstrates a lower level of anxiety and risk assessment behavior in the open field and zero maze compared to controls****. Open field A) Distance (m) traveled in the inner zone. B) Average speed (m/s) in the inner zone (PVcre 0.19 m/s ± 0.07, AcanKO 0.13 m/s ± 0.05, p-value = 0.04). Zero maze. C) Likelihood (%) of entering the whole body in the open zone after entering the head (PVcre 34.68% ± 5.93, AcanKO 60.24% ± 20.81, p-value = 0.002). Unpaired Student's t-test. Data shown as mean ± SD. Each dot represents one animal. PVcre (green) n= 10, AcanKO (purple) n = 11.*

**Table S1, qPCR primers.**

| Gene name | Forward 5'- | Reverse 5'- |
|---|---|---|
| ACAN (exon 4) | GCTTGCCTACAGAACAGCGCCA | GGGGCGTGTGGATGGGGTATCT |
| ACAN (exon 10) | CAGATGGCACCCTCCGATAC | GACACACCTCGGAAGCAGAA |
| BCAN | CTCGGCGGCTATGAGCAGTGTG | CAGGCCTCTCGTGGGTTCTGGA |
| VCAN | TGGCCCAGAACGGAAATATCA | ACTAGCCCGGAGTTTGACCAT |
| NCAN | ACGCCTACTGCTTCCGAGCTCA | GGAGGCCCCTCTGCTGACACAA |
| TNR | CCTTGCTGCGAGACCAGTGCAA | TAAAGTTGCCATGGCCGCTGCA |
| OTX2 | CGCCTTACGCAGTCAATGGGCT | GCTGCAGGACAAGAAGCCCAGG |
| SEMA3A | GGCTGGTTCACTGGGATTG | CCGTTTGCATAGTTTGCTCTGG |
| PV | GGGGACGGCAAGATTGGGGTTG | ACTGAGATGGGGCGTTGGGGAT |
| GAPDH | AGGTCGGTGTGAACGGATTTG | TGTAGACCATGTAGTTGAGGTCA |
| PPIA | TCCGACTGTGGACAGCTCTA | ATTGCGAGCAGATGGGGTAG |
